# Multi-Object Tracking in Heterogeneous environments (MOTHe) for animal video recordings

**DOI:** 10.1101/2020.01.10.899989

**Authors:** Akanksha Rathore, Ananth Sharma, Nitika Sharma, Colin J. Torney, Vishwesha Guttal

## Abstract

1. Video recordings of animals are used for many areas of research such as collective movement, animal space-use, animal censuses and behavioural neuroscience. They provide us with behavioural data at scales and resolutions not possible with manual observations. Many automated methods are being developed to extract data from these high-resolution videos. However, the task of animal detection and tracking for videos taken in natural settings remains challenging due to heterogeneous environments.
2. We present an open-source end-to-end pipeline called *Multi-Object Tracking in Heterogenous environments (MOTHe)*, a python-based application that uses a basic convolutional neural network for object detection. MOTHe allows researchers with minimal coding experience to track multiple animals in their natural habitats. It identifies animals even when individuals are stationary or partially camouflaged.
3. MOTHe has a command-line-based interface with one command for each action, for example, finding animals in an image and tracking each individual. Parameters used by the algorithm are well described in a configuration file along with example values for different types of tracking scenario. MOTHe doesn’t require any sophisticated infrastructure and can be run on basic desktop computing units.
4. We demonstrate MOTHe on six video clips from two species in their natural habitat - wasp colonies on their nests (up to 12 individuals per colony) and antelope herds in four different types of habitats (up to 156 individuals in a herd). Using MOTHe, we are able to detect and track all individuals in these animal group videos. MOTHe’s computing time on a personal computer with 4 GB RAM and i5 processor is 5 minutes for a 30-second long ultra-HD (4K resolution) video recorded at 30 frames per second.
5. MOTHe is available as an open-source repository with a detailed user guide and demonstrations at Github (https://github.com/tee-lab/MOTHe).

## 1 Introduction

Video-recording of animals is increasingly becoming a norm in behavioural studies of space-use patterns, behavioural neuroscience, animal movement and group dynamics [1, 2]. High-resolution images from aerial photographs and videos can also be used for animal census [3, 4, 5]. This mode of observation can help us gather high-resolution spatio-temporal data at unprecedented detail and help answer a novel set of questions that were previously difficult to address. For example, we can obtain movement trajectories of animals to describe space-use patterns of animals, to infer fine-scale interactions between individuals within groups and to investigate how these local interactions scale to emergent properties of groups [6, 7, 8, 9, 10, 11, 12, 13]. To address these questions, as a first step, videos need to be converted into data - typically in the form of positions and trajectories of animals. Manually extracting this information from videos can be time-consuming, tedious and, often not feasible at all. Therefore, increasingly, automated tools are being developed to detect and track animals [14, 15, 16, 17, 18, 19, 20].

However, tools developed so far work best in controlled conditions. Tracking animals from videos recorded in natural settings poses many challenges that remain unresolved. These challenges include-variability in lighting conditions, camera vibration, disappearance and appearance of animals across video frames, and heterogeneous backgrounds. Under such conditions, existing tools which rely on traditional computer vision techniques – e.g. image subtraction, colour thresholding, feature mapping, etc. – don’t perform well. In the image subtraction method [21], the motion of individuals is tracked based on differences between pixel values of two frames; this method is prone to false-detection if the camera moves, objects other than animals of interest (e.g. grass) move or if the animals don’t move. The color thresholding method [22] identifies animals in images based on their difference in colour from their background. For this method to be efficient, a consistent color/intensity difference between animals and background is necessary; this is seldom the case in natural settings because of variability in lighting conditions over both space and time and presence of other objects in the scene. Likewise, manual features (e.g. shape, orientation, edges, etc) extraction - which can be considered a generalisation of the colour thresholding - too requires consistent attributes of animals in relation to their background; consequently, this method is also likely to fail in the wild. Therefore, many popular object detection tools in ecology that use the above computer vision algorithms, although efficient for videos taken under controlled conditions, are likely to fail to detect or track animals in natural settings [23, 17].

To resolve this problem, we implement a deep learning approach. One technique found to be efficient in solving detection problems in heterogeneous backgrounds is the use of *Convolutional Neural Networks* (CNN) [24, 25, 26, 27, 28]. Neural network-based algorithms are designed based on the principles of how neurons in the visual cortex process inputs from the environment and produce an output in terms of object classification. CNNs are used to classify (or assign) categories to an image or objects within images. In the context of object detection, parts of an image are passed to the network and the network assigns a category to this image. This goal can be achieved using different approaches such as sliding window, region proposals, single-shot detector. For the classification task (i.e. assigning a category), the network uses a training dataset to learn how to classify images (i.e. sets of input pixels) to different types of output categories (e.g. animals, background, other objects of interest). The trained neural network will then be able to classify new images. Despite the promise offered by CNN-based algorithms for object detection in heterogeneous environments, only a few adaptations of them are available in the context of animal tracking [3, 29]. Recently, a few CNN-based algorithms for object detection in heterogeneous environments have been developed [30, 31, 32], but these usually require high-performance computing units such as high-end CPUs and GPUs. Additionally, implementation often requires reasonable proficiency in computer programming together with a great amount of customization. Hence, there is a need for an easily customizable end-to-end application that automates the task of object detection and is usable even on simple desktop machines.

Here, we provide an open-source package, *Multi-Object Tracking in Heterogeneous environment* (MOTHe), which is easy to customize for different datasets and can run on relatively basic desktop units. MOTHe can detect and track multiple individuals in heterogeneous backgrounds. It uses a color thresholding approach followed by a small CNN architecture to detect and classify objects within images, allowing fast training of the network even on relatively unsophisticated desktop computing units. The network can then be used on new images to detect animals. The code then generates individual tracks from detections using a Kalman filter. It provides an end-to-end pipeline that automates each step including the training data generation, detection, and tracking. In this paper, we have implemented MOTHe on six video clips from two species (wasps on the nests and antelope herds in four different types of habitats). These videos were recorded in natural and semi-natural settings having background heterogeneity and varying lighting conditions. We also provide an open to use GitHub repository (https://github.com/tee-lab/MOTHe) along with a detailed user guide for the implementation.

## 2 Working principle & features

MOTHe is a python-based library and it uses a Convolutional Neural Network (CNN) architecture for object detection. CNNs are specific types of neural network algorithms designed for image classification (assigning a category to an image or part thereof). It takes a digital image as an input and processes pixel values through a network and assigns a *category* to the image. To achieve this, CNN is trained via a large amount of labeled training data and learning algorithms; this learning procedure enables the network to learn features of objects of interest from the pool of training data. Once the CNN models are trained, these models can be used to classify new data (images). In the context of tracking multiple animals in a video, an object detection task would involve identifying locations and categories of objects present in an image. MOTHe works for 2-category classification e.g. animal and background.

In this section, we present a broad overview of features and principles on which MOTHe works. Details of all user-inputs and guidelines to run and customize the modules are available in a user manual on the Github repository and also in the supplementary material. MOTHe’s network architecture and parameters are fixed to make it user-friendly for beginners. However, advanced users can modify these parameters and tweak the architecture in the code files.

MOTHe is divided into four independent modules (see Figure 1):

i. **Generation of training dataset** - Dataset generation is a crucial step in object detection and tracking. Generating enough data for training takes a lot of time if done manually. In this step, we automate the data-generation. Users run the command line code to extract images for the two categories i.e. animal and background. It allows users to crop regions of interest by simple clicks over a Graphical User Interface and saves the images in appropriate folders. On each run, users can input the category for which the data will be generated and specify the video from which images will be cropped. Outputs from this module are saved in two separate folders one containing images of animals (yes) and the other containing background (no).
ii. **Network training** - The network training module is used to create the network and train it using the dataset generated in the previous step. Users run a command-line script to perform the training. Once training is complete, the training accuracy is displayed and the trained model (classifier) is saved in the repository. The accuracy of the classifier is dependent on how well the network is trained, which in turn depends on the quality and quantity of training data (see section “How much training data do I need?” in Supplementary Materials). Various tuning parameters of the network, for e.g; number of nodes, size of nodes, convolutional layers etc., are fixed to render the process easy for the user.
iii. **Object detection** - To perform the detection task, we first need to identify the areas in an image where the object can be found, this is called localization or region proposal. Then we classify these regions into different categories (eg whether an animal or background?), this step is called *classification*. The object detection module uses the trained CNN model and performs above two key tasks on any given input image: Localisation and classification. The localisation step is performed using an efficient thresholding approach that restricts the number of individual classifications that need to be performed on the image. The first stage grayscale thresholding will output pixels that contain animals along with false positives (i.e. the locations in the background that have a similar color profile to the animals). The classification at each location is then performed using the trained CNN generated in the previous module. The outputs, detected animals, are in the form of .*csv* files that contains locations of identified animals in each frame.
iv. **Track linking** - This module assigns unique IDs to the detected individuals and generates their trajectories. We have separated detection and tracking modules so that the package can also be used by someone interested only in the count data (eg. surveys). This modularisation also provides flexibility by allowing the use of more sophisticated tracking algorithms to experienced users. We use a standard approach for track linking that uses a Kalman filter to predict the next location of the object and the Hungarian algorithm to match objects across frames. This script can be run once the detection output is generated in the previous step. Output is a .*csv* file that contains individual IDs and locations in each frame. Video output with unique IDs on each individual is also generated.

**Figure 1:**
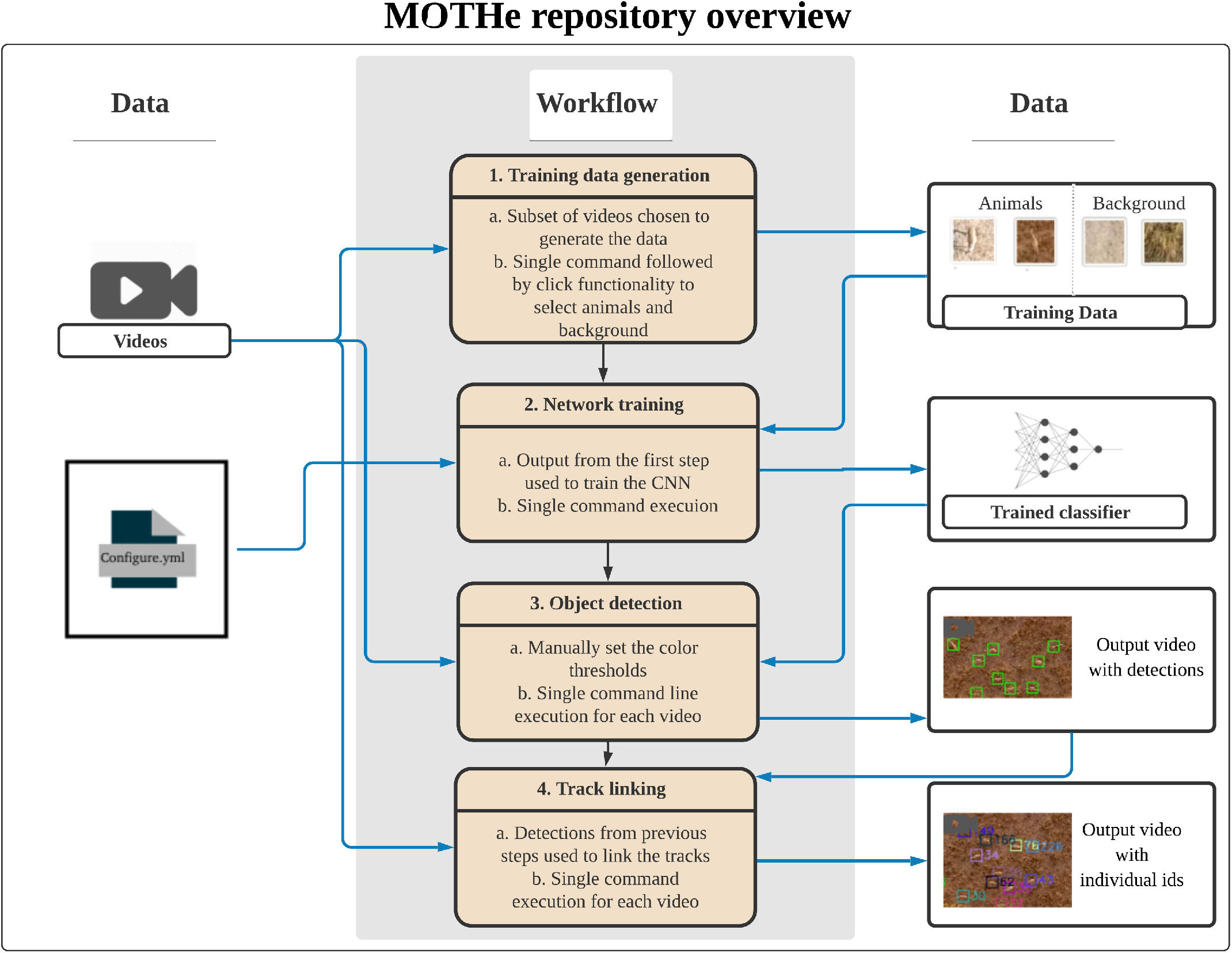
The layout of our Github repository. A configuration file is generated in the first step, which maintains directory paths and parameter values used by subsequent modules. Tracking happens in two steps-first, we need to train the network on training dataset; second, object detection is done usingthe trained CNN on the image. Each step here is a separate module that can be run by users. Black arrows represent the directional flow of executable files. Blue arrows represent input/output flow of data in the modules.

### 2.1 Localisation and compact network

Two features make MOTHe fast to train and run on new videos - localisation using grayscale-thresholding approach and a compact CNN architecture. In MOTHe, to achieve localisation, we use threshold on the grayscale image to identify key-points where an animal may be located. This step reduces the computation time compared to a sliding window approach [33]. To further reduce the computation time, we have used a compact architecture with only six convolutional layers. The use of a compact CNN architecture also has the advantage of requiring smaller training datasets and is less prone to overfitting than deeper networks.

A trade-off of the above two approaches is the reduced generality of the trained model across different types of datasets. To deal with this drawback, we provide options to change parameters for different datasets so that the network retains its accuracy for a specific detection task. We demonstrate our software pipeline on two different video datasets which are explained in the next section.

## 3 Implementation on example videos

To demonstrate our the application of repository,, we used videos of two species - blackbuck (*Antilope cervi-capra*)and a tropical paper wasp (*Ropalidia marginata*). These two species present varying complexity in terms of the environment (natural and semi-natural settings), background, animal speed, behaviour and overlaps between individuals (Figure 2). Below, we provide a description of these datasets and describe the steps to implement MOTHe (see Figure 1 for overview).

**Figure 2:**
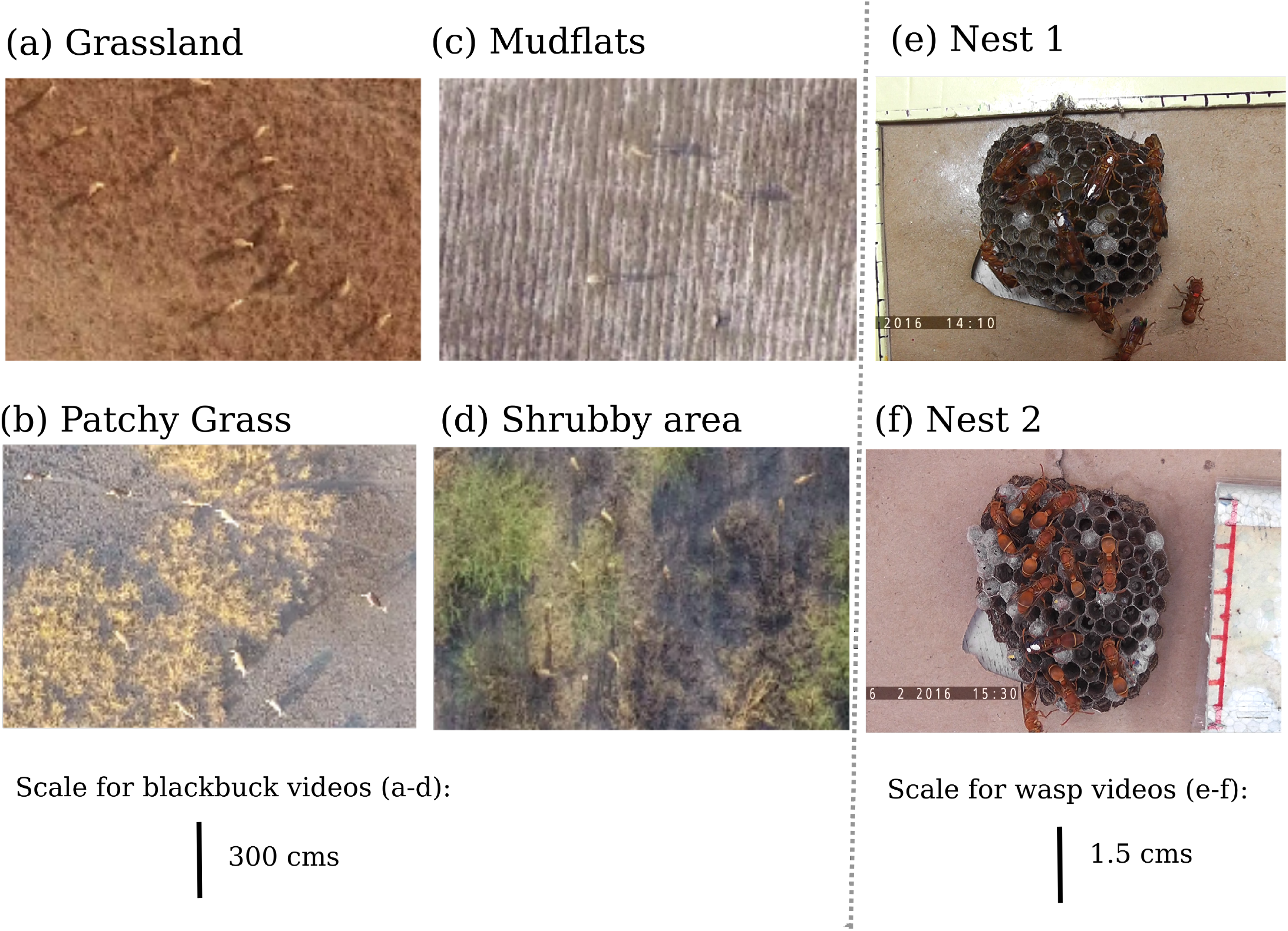
Variation in the appearance of animals and background in different videos. Blackbuck herds in a (a) grassland, (b) habitat having patches of grass, (c) mudflat area of the park, (d) bush dominated habitat. Wasp nest with a majority of (e) older wasps, (f) newly eclosed wasps.

### 3.1 Data description

#### 3.1.1 Collective behaviour of blackbuck herds

We recorded blackbuck (*Antilope cervicapra*) group behaviour in their natural habitat. Blackbuck herds exhibit frequent merge-split events [34]. These herds consist of adult males & females, sub-adults and juveniles [35, 36]. They are sexually dimorphic and the colour of adult males also changes with testosterone levels [37]. This colour variation makes it difficult to use color segmentation based techniques to detect them. Major source of complexity in this system arises from their heterogeneous habitat, comprising of semi-arid grasslands with patches of trees and shrubs. While many blackbuck don’t move across many video frames, there is a substantial movement of grasses and shrubs in the background. These conditions pose challenges for applying basic computer vision methods such as colour thresholding and image subtraction.

We recorded blackbuck movement in different habitat patches - grasslands, shrublands, and mudflats. These recordings were collected using a DJI quadcopter flown at a height of 40-45 meters (Phantom Pro 4) equipped with a high-resolution camera (4K resolution at 30 frames per second). The average size of an adult blackbuck is 120 cms from head to tail which corresponds to around 35 pixels in our videos.

#### 3.1.2 Nest space-use by wasps

We used videos of a tropical paper wasp *Ropalidia marginata* recorded under semi-natural conditions [38]. Here, individuals were maintained in their natural nests in laboratory conditions and were allowed to forage freely.

Nests of *Ropalidia marginata* are sites for social interactions between mobile adults as well as between adults and immobile brood [39]. These nests are made of paper, which offers a low contrast to the dark-bodied social insects on the nest surface. Nest comprises of cells in which various stages of brood are housed and thus add to the heterogeneity of the background. Additionally, different nest colonies differ in the age composition of individuals, contributing to the variation in the appearance of wasps across videos. Therefore, this system too presents challenges to classical computer vision methods used to detect animals from the background. Recordings were done using a video camera (25 frames per second). The size of wasp is 1 cm from head to the abdomen which corresponds to around 150 pixels in our videos.

### 3.2 Parametrisation

For the ease of use, we have kept the parameterisation process minimal. Therefore, the various tuning parameters of the network architecture are fixed. However, advanced users can change the parameters in the code to customise it for more sophisticated tasks. The only step which requires some amount of parameter scanning by users is choosing the *color thresholds* for animals. As mentioned earlier, to improve speed of processing, we use a color thresholding approach as the localisation step. For this technique, users need to input values of the minimum and maximum threshold of the pixel values that may potentially correspond to the animal; these numbers are to be edited in the *config.yml* file. To choose the values for any generic dataset, we provide detailed instructions under the section *Choosing color threshold* of the Github repository.

### 3.3 Data generation and CNN Training

To generate data for training the CNN, we run a simple one-line command which then displays frames from the videos; for each of these frames, we select animals and background examples that are used in the training step. The resulting output is automatically stored in separate folders for animals and backgrounds (see *Using MOTHe, Step 2* in the Github repository).

To generate training samples for blackbuck videos, we used 2000 frames from 45 different videos; these videos were from different types of habitats. The number of individuals in these videos ranges from 30-300 individuals. We fixed the values of the gray-scale threshold for blackbuck to be [0,150]. We then run the CNN training command (see *Using MOTHe*, Step 3 in the Github repository).

Likewise, to generate data and train the network with features of wasps we used equally spaced 1000 frames from 6 different videos. We fixed the values of the gray-scale threshold for wasps to be [150,250].

### 3.4 Detection and track-linking

We now present results after running the trained CNN on four sample videos of blackbuck herds, representing different habitat types and the group sizes (Figure 2 (a)-(d)) and two sample videos of wasps, representing two different colonies (Figure 2 (e)-(f)). The sample videos were all 30 seconds long. The maximum number of individuals present in these videos are 156 and 12 for blackbuck and wasps, respectively.

In Figure 3, the first column shows the results of running object detection on these video clips and the second column displays the results after implementing track linking on the detections. We observe that the package is able to detect and track nearly all individuals in all types of habitat. However, as expected, there are some errors in animal detection using MOTHe.

**Figure 3:**
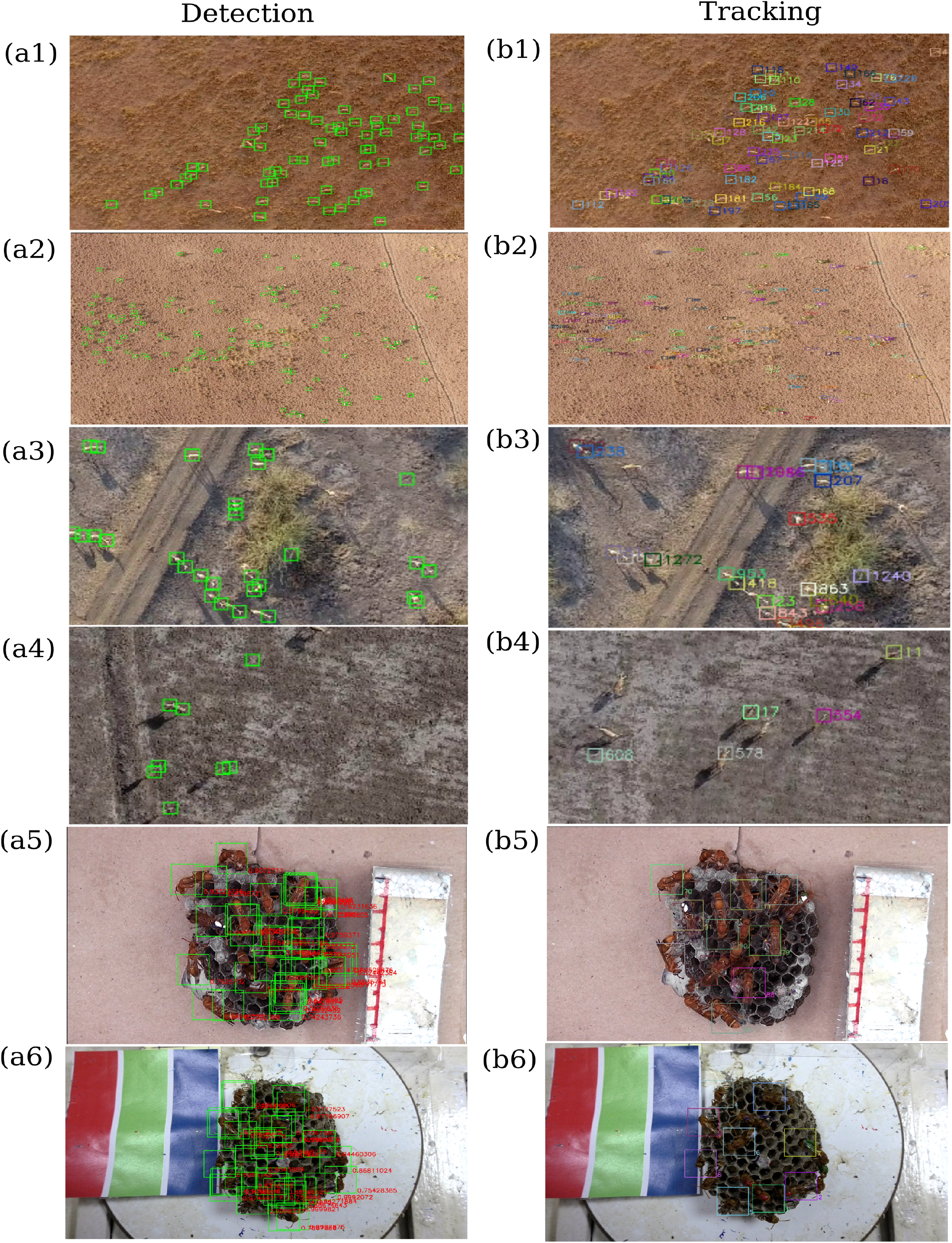
Detection and Tracking results in six example videos. (a1) and (b1) Moderate size blackbuck herd in a grassland; (a2) and (b2) A big herd (blackbuck - 158 individuals) in the grassland; (a3) and (b3) blackbuck herd in a shrubby area; (a4) and (b4) blackbuck herd in the mudflats; (a5) and (b5) Nest with a majority of older wasps and (a6) and (b6) Nest with a majority of newly eclosed wasps. All images are zoomed and scales at different levels for visibility. The size of wasps is around 1 cm and blackbuck is around 1 meter.

To quantify these error rates in MOTHe, we prepared *ground-truth* data by visually counting the number of individuals present in each frame and compared it with the number of detections obtained by running MOTHe; this was repeated for 30 frames spaced at 1 second and for each of the videos. This quantification gives us ground-truth values of animals and detections. We then compared this with the detections of animals using the MOTHe on the same set of frames to obtain (i) the percentage true detections and (ii) percentage false detections (i.e. arising from the wrong classification of background objects as animals). The results, shown in Table 1, demonstrate fairly high true detection rates (of 80% and above) and low false detection rates (close to zero in most videos).

**Table 1:** Results after running MOTHe on blackbuck videos in various habitat and wasp videos in two colonies. Each video clip is of 30 seconds in duration and these results are averaged over 30 frames spaced at 1 second for each video. % true detections show the number of individuals that were correctly detected in a frame and % false positives show the background noise identified as an animal. For computing efficiency, run-time in frames processed per second is reported.

| Video | Group size | Habitat | % true detections | % false detections | Run time (Frames processed per sec.) |
| --- | --- | --- | --- | --- | --- |
| Blackbuck-1 | 28 | Patchy grass | 89.3 | 14.2 | 1.99 |
| Blackbuck-2 | 78 | Grass | 83.1 | 0 | 0.82 |
| Blackbuck-3 | 156 | Grass | 97.4 | 0.64 | 0.51 |
| Blackbuck-4 | 34 | Shrubs | 91.4 | 0 | 2.44 |
| Wasp-1 | 15 | Colony with majority older wasps | 86.6 | 0 | 1.11 |
| Wasp-2 | 16 | Colony with newly eclosed wasps | 93.75 | 0 | 1.06 |

**Table 2:** Comparison of MOTHe with other popular tracking solutions in terms of three qualitative features: video complexity, ease of use and performance. * Performance quantified by running these techniques on blackbuck videos, all other run-time are as reported by the authors.

|  | Features | MOTHe | Tracktor | idTracker | Yolo v3 | BioTracker | ToxTrac |
| --- | --- | --- | --- | --- | --- | --- | --- |
| Video Complexity | Detection against heterogeneous background? | yes | for single individual | no | yes | no | no |
|  | Multiple individual tracking? | yes | in homogeneous background | yes | yes | yes | yes |
|  | Identifies stationary animals as well? | yes | yes | yes | yes | no | NA |
| Ease of use | Requires sophisticated infrastructure? (GPUs) | no | no | minimum 8GB RAM | yes | no | no |
|  | Interface and installation | Command based | Command based | GUI | Command based | GUI | GUI |
|  | Click and drag functionality for training-data generation | yes | NA | NA | no | NA | NA |
| Performance | Maximum number of individuals tracked in test run | 156 | 8 | 35 | - | 11 in the example figure | 20 |
|  | Computational efficiency | 180 frames/minute for 4K resolution video* | 9 minutes 43 seconds for 33 MB video (fish schooling) | 2s per frame for a HD video with 20 medaka fish | 5 frames/minute for 4K resolution video* | NA | 25 frames per second in HD videos using modern computers |
|  | Species on which testing was done | Antelope, Wasp | Fish, spider, termite, mice, tadpole | Fish, ant, mice, flies | Wildebeest, Zebra | Fish | Fish, mice, cockroach, ant |
|  | Tested in conditions | Natural and semi-natural | Controlled environment for multiple individuals | Common lab conditions and manipulations | Natural | Controlled | Controlled |

We emphasise that even if some animals were not detected in particular frames, they were detected in the subsequent frames. Therefore, all the wasps and blackbuck present in our video clips were tracked by MOTHe (see Supplementary Videos).

We also show the time taken to run detection on these video clips (Table 1) on an ordinary laptop (4 GB RAM with an Intel Core i5 processor); we find that the number of frames processed in one second ranged from 0.5 to 2.5. This efficiency can be improved considerably by running MOTHe on workstations, GPUs or cloud services.

## 4 Discussion

We demonstrate that MOTHe is relatively easy to use software pipeline that allows users to generate datasets, train a simple neural network and use that to detect multiple objects of interest in the heterogeneous background. We demonstrated the application of MOTHe on two relatively different types of systems in which the animal species, their movement type, animal-background contrast, and background heterogeneity all differ. We argue that MOTHe is potentially applicable to a wide variety of animal videos in their natural conditions.

The use of machine learning for classification enables MOTHe to detect stationary objects. This bypasses the necessity of relying on the motion of animals for the detection of animals [15]. MOTHe has various built-in functions and is designed to be user-friendly; advanced users can customize the code to improve the efficiency further. MOTHe is modular, organized and (semi-)automated which helps the user to achieve object tracking with relatively minimum efforts. MOTHe can be used to track objects on a desktop computer or a basic laptop. Alternative methods for object detection, such as YOLO [31] or RCNN [28, 27] that perform both localisation and classification, are expected to reduce error rates compared to our approach and do not require colour thresholding. However, these types of neural network require access to high specification GPUs. Using these kinds of specialised object detectors for animal tracking requires sufficient user proficiency to configure. Furthermore, these methods are not typically tailored to the detection of small objects in high-resolution images.

MOTHe performs well even in scenarios with poor object contrast with the background, bad lighting, background noise, and viewpoint. The use of CNN in this package accounts for morphological variations and scaling issues. The use of machine learning algorithms makes MOTHe highly versatile and training the CNN with sufficient sample images results in high accuracy for detection in complex settings. However, like most tracking algorithms [14, 15, 16, 17, 18, 19], MOTHe is incapable of resolving tracks of individuals in close proximity (usually, when less than one body length). As a trade-off to its computational efficiency, we did not incorporate issues arising from a shaking camera in the MOTHe application. Nonetheless, it can be used in combination with image stabilizing algorithms to solve camera shake issues or could be resolved by smoothing the trajectories after processing.

In our examples, the maximum number of individuals presented to the detection algorithm was 156 and MOTHe was able to detect all 156 individuals. We surmise that it should be able to detect even larger numbers of individuals as long as the distance between individuals is at least one body length. In table 3, we compare MOTHe’s qualitative performance with some popular tracking solutions. In future studies, a detailed quantitative comparison of several computer-vision based object detection techniques on different types of datasets could be useful for researchers to choose among many options available.

In summary, MOTHe allows researchers with relatively minimal coding experience to track stationary as well as moving animals in their natural habitats. For each step of the detection and tracking process, users need to run a single command. MOTHe is available as an open-source repository with a detailed user guide and demonstrations via Github. We believe that this end-to-end package will encourage more researchers to use video observations for studying animal group behaviour in natural habitats and would be of use to a larger research community.

## Acknowledgments

We are grateful to Ashwin Karichannavar for testing the MOTHe repository and providing inputs. We also thank Hari Sridhar, Vivek Hari Sridhar and Hemal Naik for providing critical feedback on the manuscript. VG acknowledges support from DBT-IISc partnership program and infrastructure support from DST-FIST. AR and NS thank MHRD for the Ph.D. scholarship. We acknowledge a UGC-UKIERI for a collaborative research grant between VG and CJT. We thank the forest department of Gujarat for the logistical support and the permission to work in Blackbuck National Park, Velavadar. The associated animal research was approved by the Institutional Animal Ethics Committee at the Indian Institute of Science.

## Author Contributions

AR conceptualized the project with inputs from VG. AR, AS and CJT contributed methods. AR and NS contributed data. AR, AS and NS analysed the data. AR and VG synthesized the results and wrote the paper with inputs from coauthors.

