## Supplementary Material for "Multi-Object Tracking in Heterogeneous environments (MOTHe) for animal video recordings"

### MOTHe

Mothe is a pipeline developed to detect and track multiple animals in a heterogeneous environment. MOTHe is a python based repository and it uses *Convolutional Neural Network(CNN)* architecture for the object detection task. It takes a digital image as an input and reads its features to assign a category. These algorithms are learning algorithms which means that they extract features from the images by using huge amounts of labeled training data. Once the CNN models are trained, these models can be used to classify novel data (images). MOTHe is designed to be generic which empowers the user to track objects of interest even in a natural setting.

**MOTHe has been developed and tested on Ubuntu 16.04 using python 3.5.2**

### PIPELINE DESCRIPTION:

MOTHe can automate all the tasks associated with object classification and is divided into 5 executables dedicated to the following tasks.

1. **System configuration**: The system configuration is used to setup MOTHe on the users system. Basic details such as the path to the local repository, path to the video to be processed, the size of the individial to be cropped and the size of the bounding box to be drawn during the detection phase.

2. **Dataset generation**: The dataset generation is a crucial step towards object detection and tracking. The manual effort required to generate the required amount of training data is huge. The data generation class and executable highly automates the process by allowing the user to crop the region of interest by simple clicks over a GUI and automatically saves the images in appropriate folders.

3. **Training the convolutional neural network**: After generating sufficient number of training example, the data is used to train the neural network. The neural network produces a classifier as the output. The accuracy of the classifier is dependent on how well the network is trainied, which in turn depends on the quality and quantity of training data (See section “**How much training data do I need?**”). The various tuning parameters of the network are fixed to render the process easy for the users. This network works well for binary classification - object of interest

(animals) and background. Multi-class classification is not supported on this pipeline.

4. **Object detection**: This is the most crucial module in the repository. It performs two key tasks - it first identifies the regions in the image which can potentially have animals, this is called localisation; then it performs classification on the cropped regions. This classification is done using a small CNN (6 convolutional layers). Output is in the form of *.csv* files which contains the locations of the identified animals in each frame.

5. **Object tracking**: Object tracking is the final goal of the MOTHe. This module assigns unique IDs to the detected individuals and generates their trajectories. We have separated detection and tracking modules, so that it can also be used by someone interested only in the count data (eg. surveys). This modularisation also provides flexibility of using more sophisticated tracking algorithms to the experienced programmers. We use an existing code for the tracking task (from the Github page of ref). This algorithm uses Kalman filters and Hungarian algorithm. This script can be run once the detections are generated in the previous step. Output is a \textit{.csv} file which contains individual IDs and locations for each frame. A video output with the unique IDs on each individual is also generated.

**MOTHe flowchart**

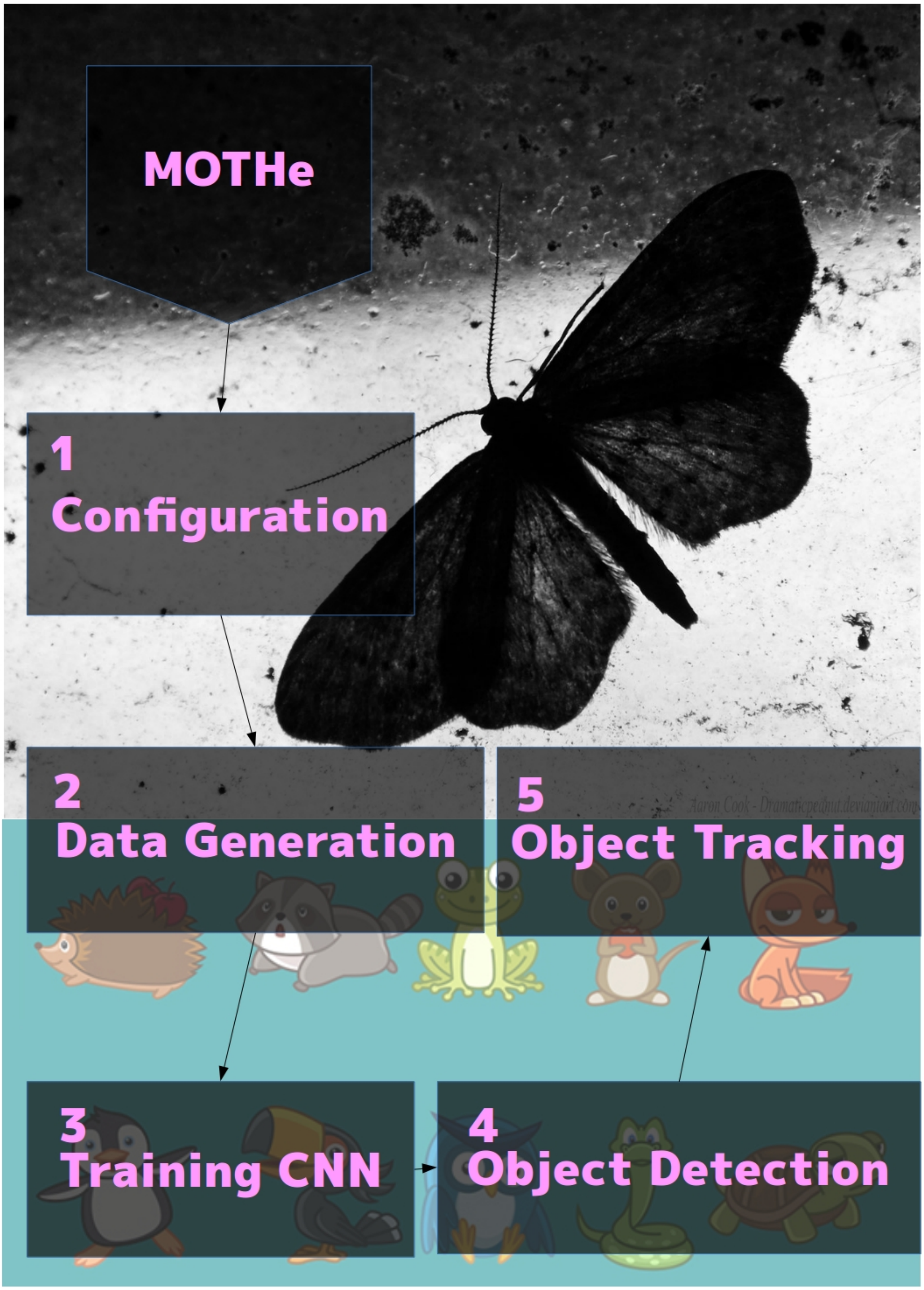


The flowchart depicts the order of the functions to be used in MOTHe. The user should follow the following steps to successfully detect and track multiple animals using MOTHe

### INSTALLATION

The current version of MOTHe is supported for Linux (ubuntu) machines. We provide command line user functionality for all the executable files of MOTHe. The usage of MOTHe requires certain packages to be installed on your ubuntu machine. A *mothe.sh* shell script is provided which helps in the installation of these packages in a single step.

To get started with MOTHe, you need to download the MOTHe git repository on your local machine. Open the terminal and use the 'git clone' command to clone the MOTHe repository as shown in the figure below. You can also download the MOTHe repository directly from the github page.

```
$ git clone https://github.com/tee-lab/MOTHe/
```

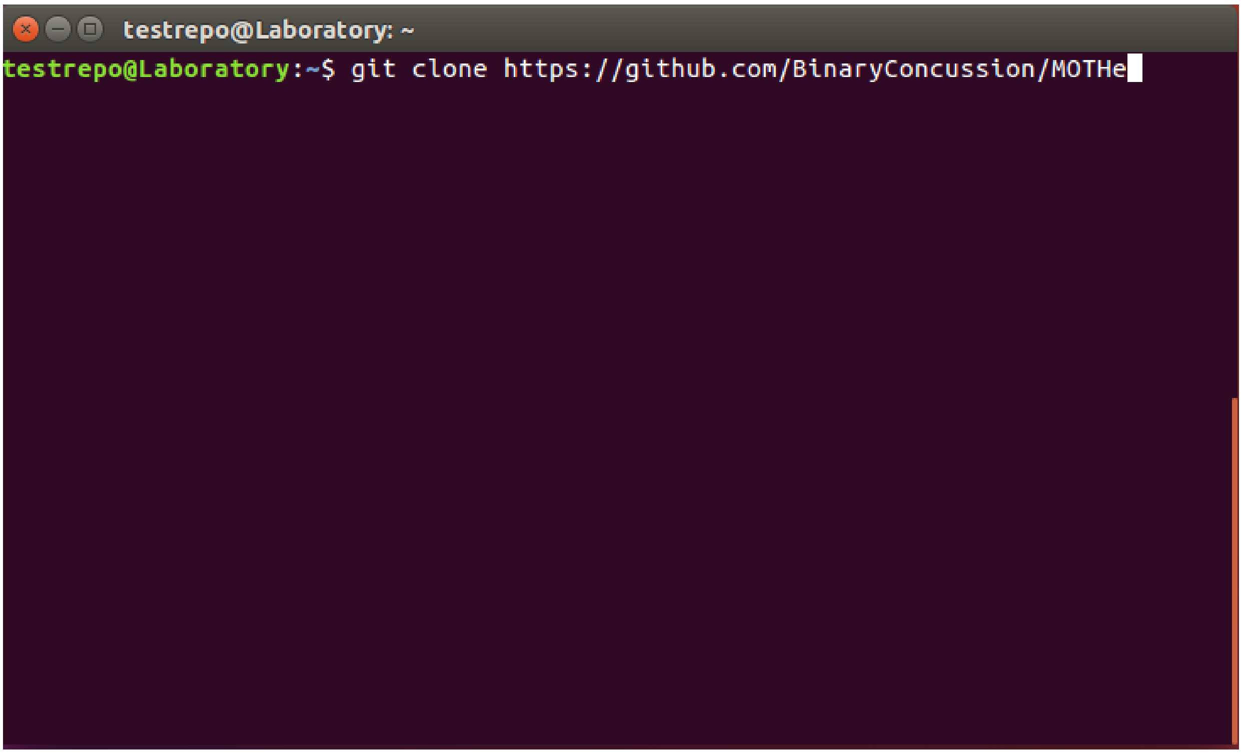


Once the repository is succesfully clones, you will see below output on the screen.

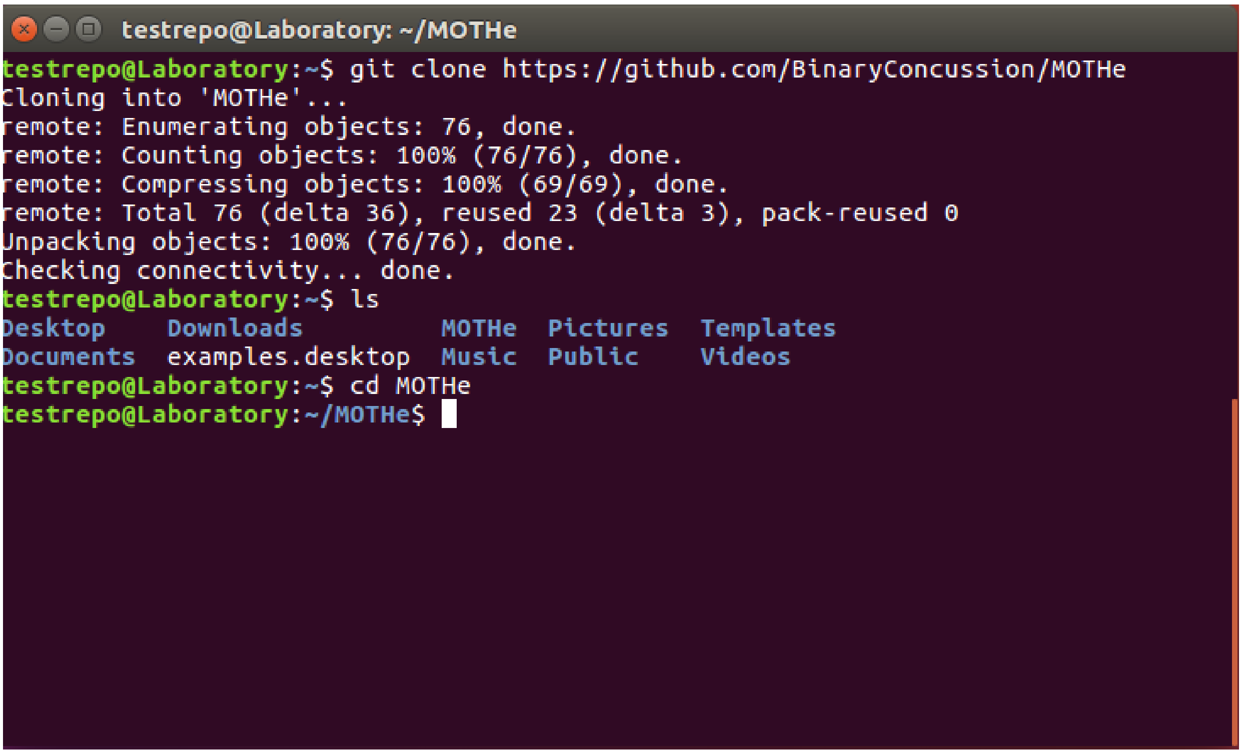


After downloading the repository on the local system, open the terminal and navigate to the local MOTHe directory as shown below:

```
$ cd Desktop/user/MOTHe
```

Replace above path with the path to MOTHe repository in your system.

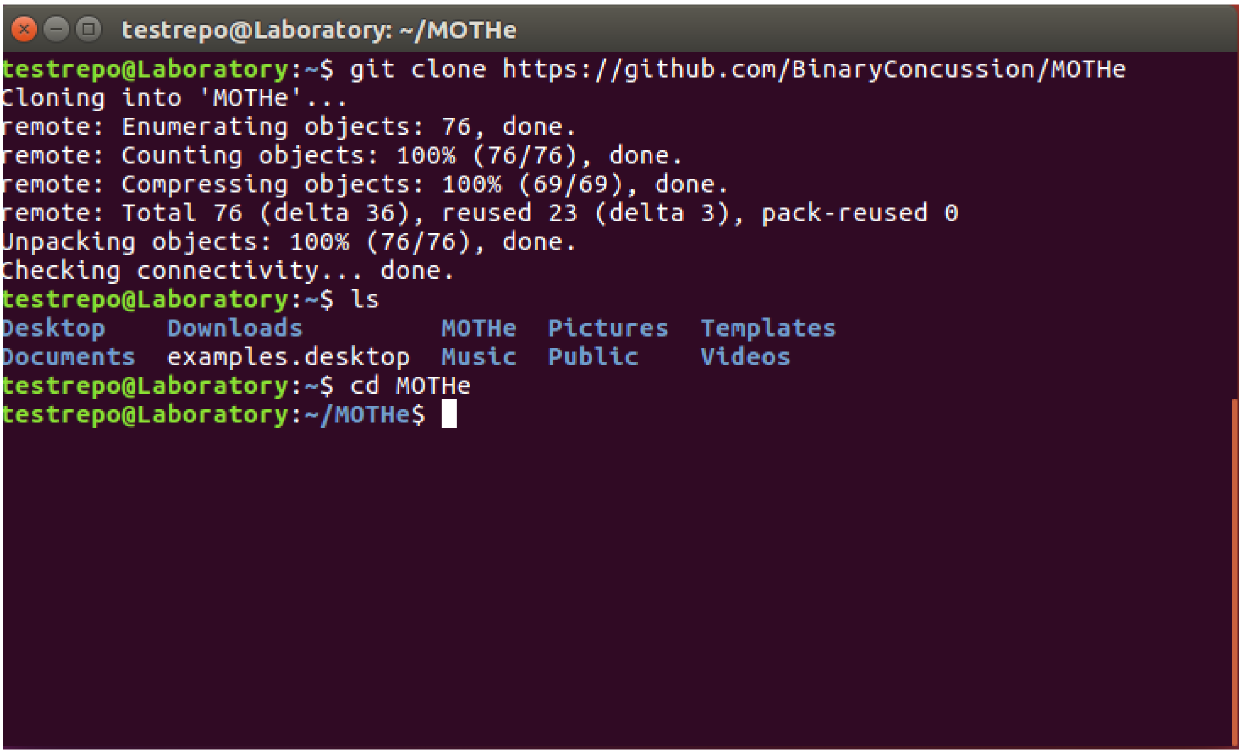


The usage of MOTHe requires certain packages to be installed on your ubuntu machine. A [mothe.sh] shell script is provided which helps in the installations of these packages in a single step. List the files in the MOTHe directory and take note of the [mothe.sh] file

```
$ ls
```

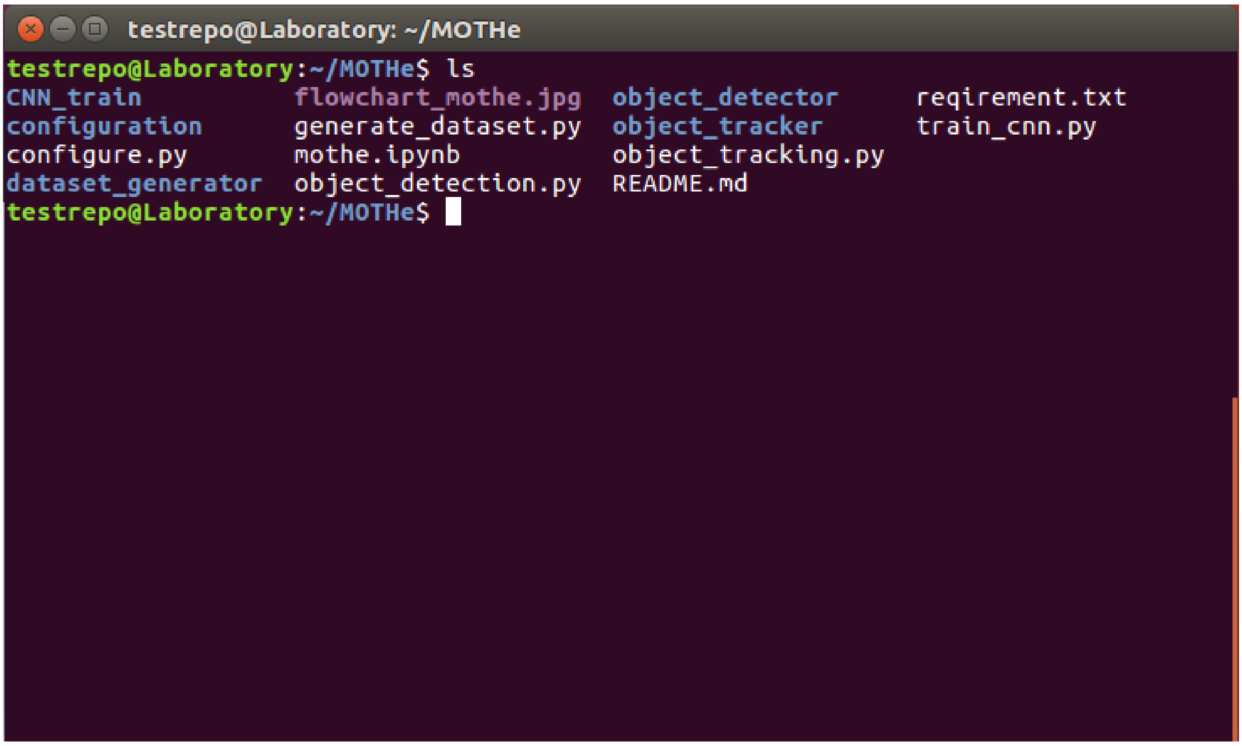


To use the [mothe.sh](http://mothe.sh) script, you have to first change the permissions to the file using the following command

```
$ chmod +x ./mothe.sh
```

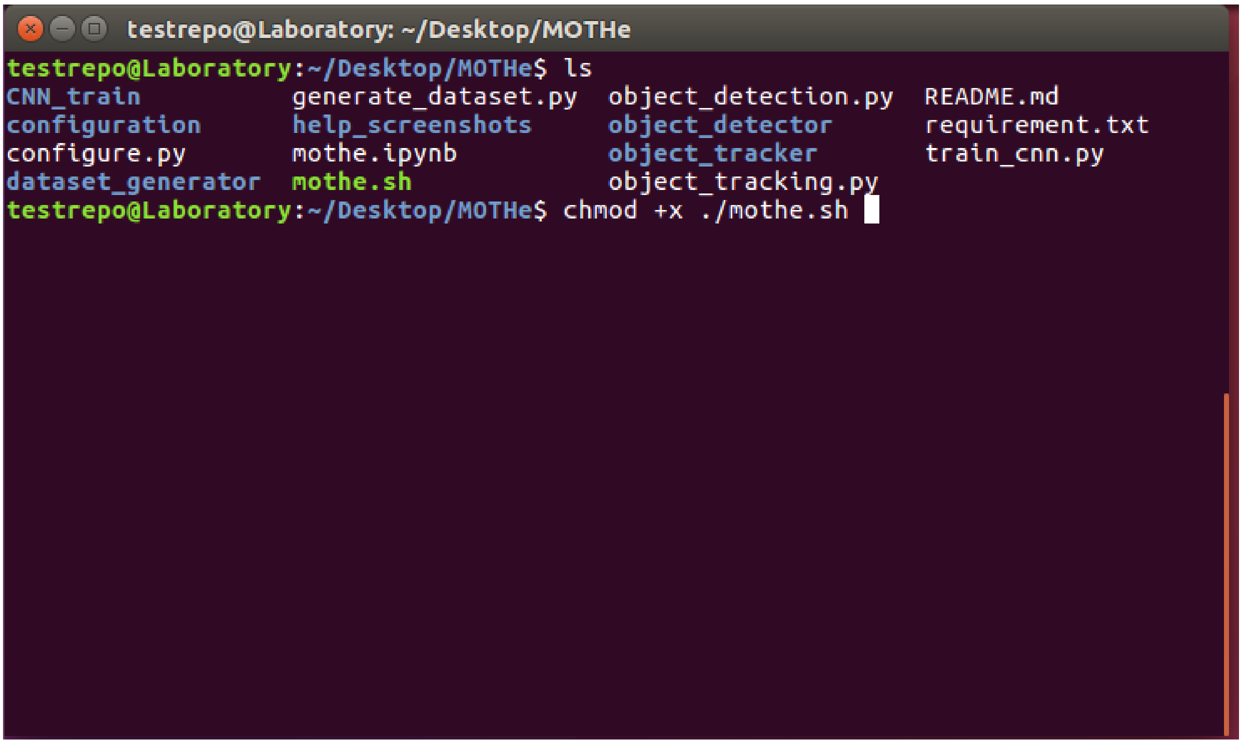


After changing the permissions to the [mothe.sh](http://mothe.sh) file, use the bash command to use the shell script as shown below

```
$ sudo bash mothe.sh
```

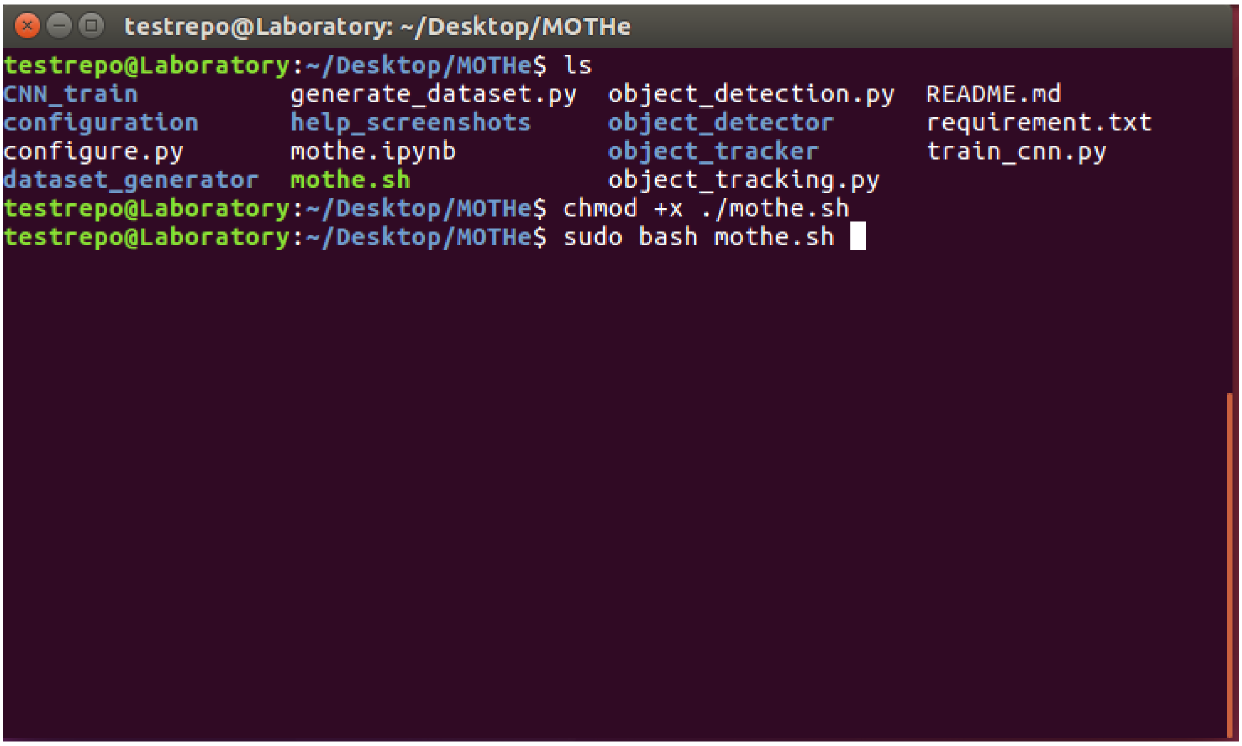


The *mothe.sh* also installs and configures a virtual environment where an environment can be created specific to MOTHe. Create a virtual environment by the name of mothe as shown below

```
$ mkvirtualenv mothe -p python3
```

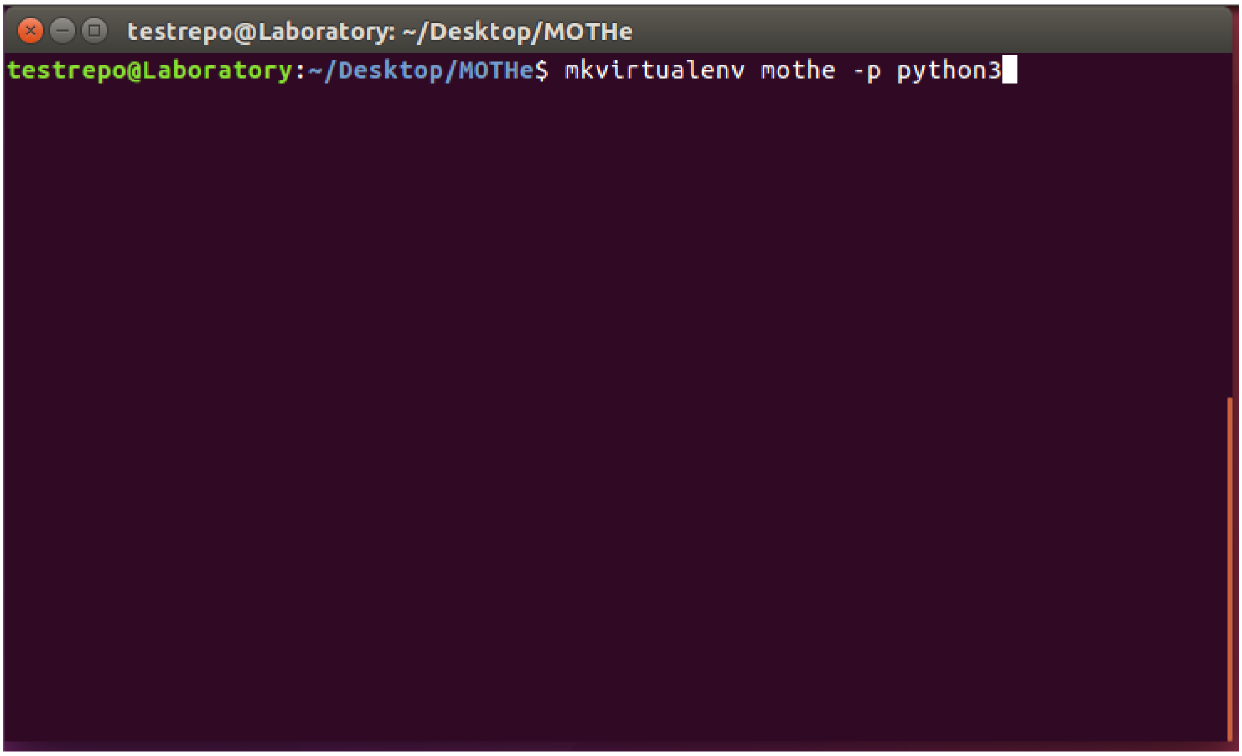


After creating the the virtual environment, activate the virtual environment as follows

```
$ workon mothe
```

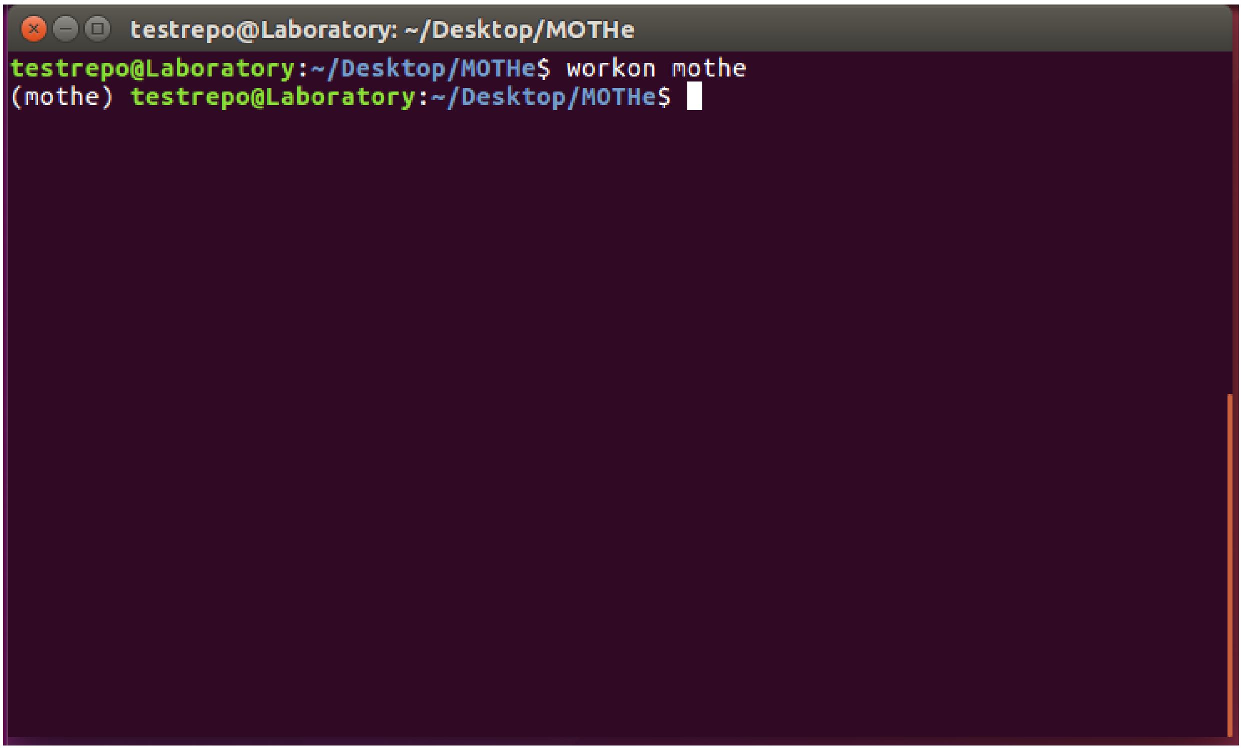


After activating the mothe environment, configure all python modules using the *requirement.txt* as follows:

```
$ python3 -m pip install -r requirement.txt
```

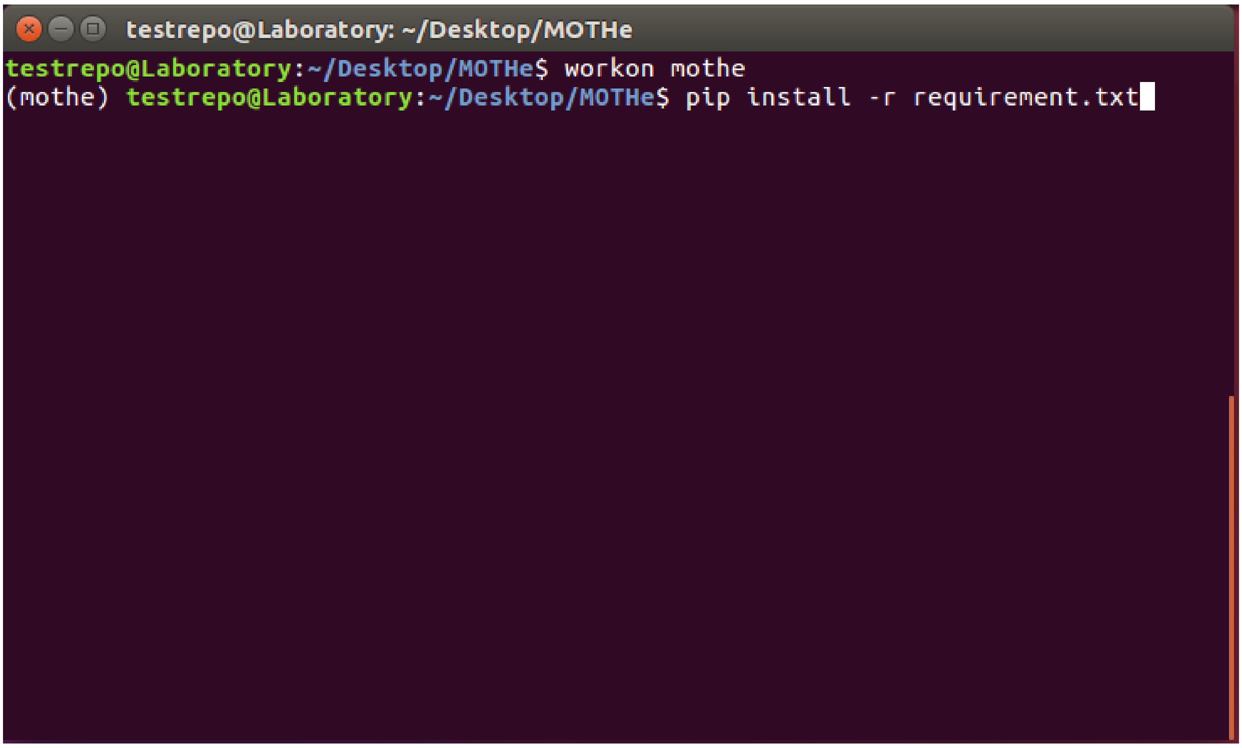


User also has an option to install these modules manually. Open the *requirement.txt* file and install all the modules listed in this file.

### USING MOTHe

Users can run MOTHe to detect and track single or multiple individuals in the videos (or images). In this section, we describe the step-by-step procedure to run/test MOTHe. If you are interested in learning the procedure first by running it on our videos (and data), follow the guidelines under subsection **"Testing"** in each step.

1. **Step 1 - System configuration**: File name - *configure.py*

This step is used to set parameters of MOTHe. All the parameters are saved in *config.yml*. Parameters to be set in this step - home directory, cropping size of animals in the videos, path to video files etc. It wil also prompt to enter runmode, if you are exploring MOTHe on blackbuck or wasp data, enter 0 otherwise for your data enter 1.

Run the following command in your terminal from your MOTHe directory-

```
$ python configure.py
```

Once the above command is executed, user will be prompted to enter path to MOTHe directory in user's system.

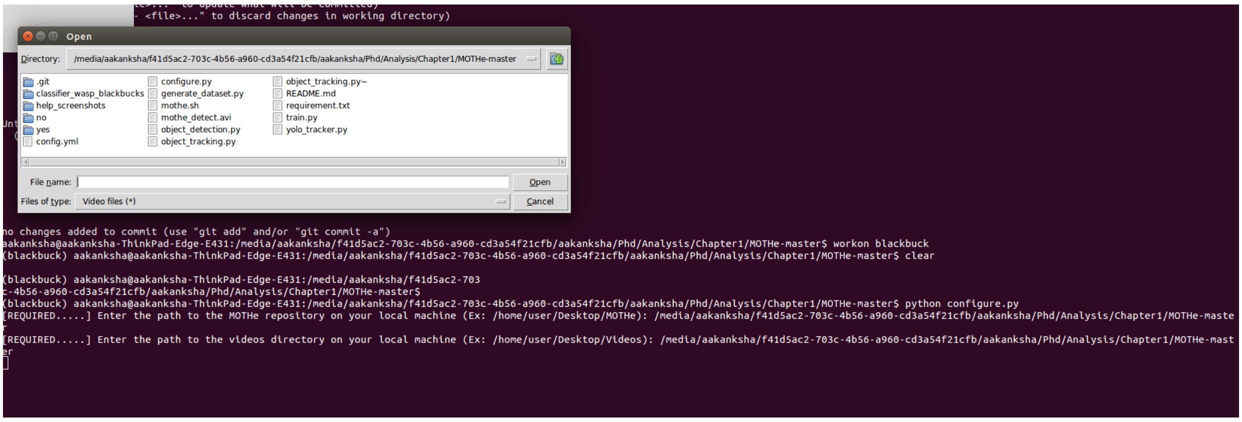


After entering the path, a prompt for selecting a video will appear. Select any video from the videos in which you want to track the animals and draw a bounding box around the animal. Press **c** to confirm the selection, **r** to discard and draw again.

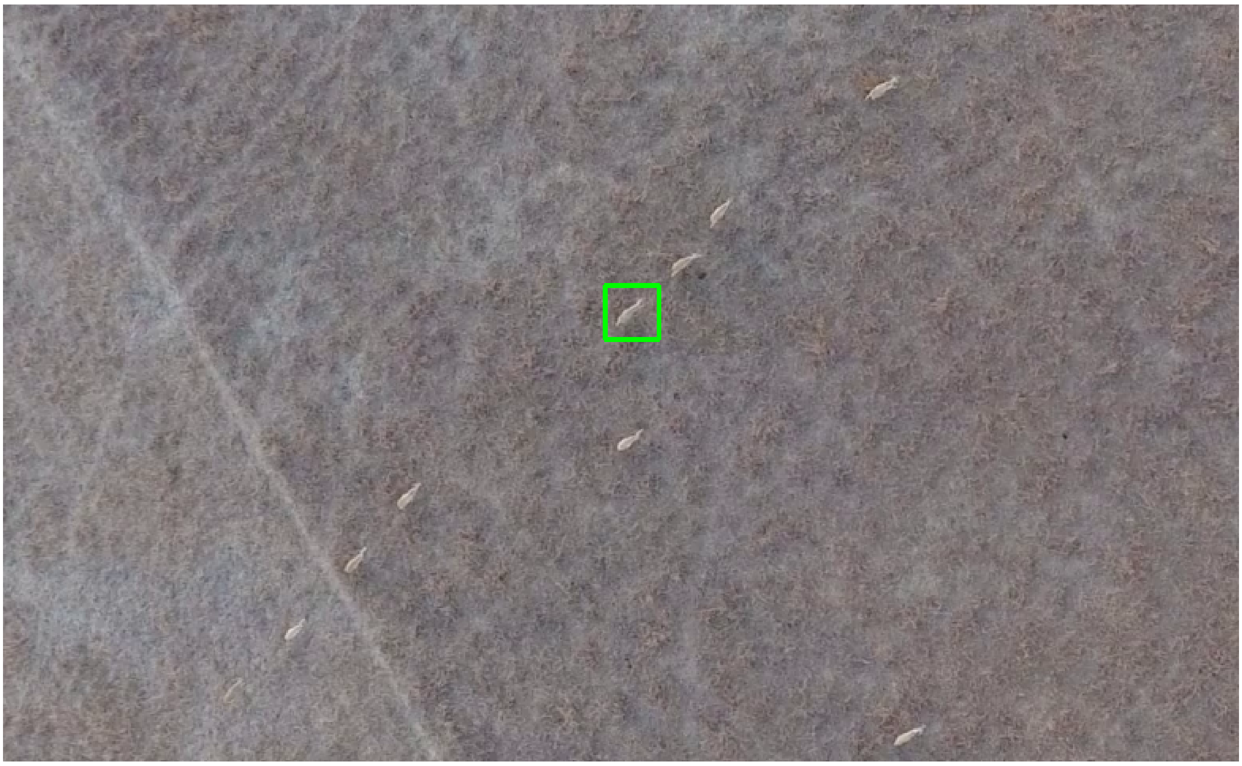

2. **Step 2 - Dataset generation**: File name - *generate_dataset.py*

This program will pick frames from the videos, user can click on animals or background in these images to create samples for both categories (animal and background). Examples of animal of ineterst will be saved in the folder **yes** and background in the folder **no**. User needs to generate at least 8k-10k samples for each category (see section **How much training data do I need?** for detailed guidelines). One must ensure to take a wide representation of forms in which animals appears in the videos and same for the background variation.

To generate the training data, run the following code in the terminal:

```
$ python generate_dataset.py {step}
```

Replace `{step}` with the number of frames you want to skip. This number will depend on the duration and number of videos. For example, if you have 20 10-minute videos at 30 FPS and on an average 50 animals in each frame, to generate 10k images we need at least 500 images from each video. On an average we will get 50 animals from one frame so we need 10 frames from each video. So, in this case we would set step size as 150.

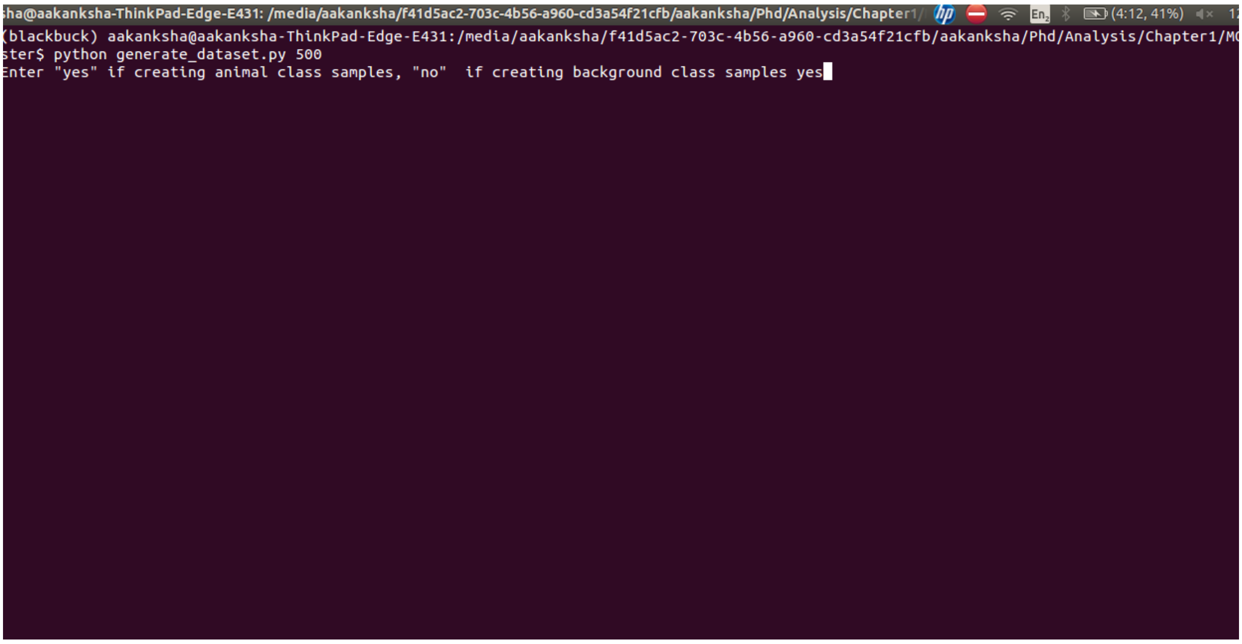


Once the code is run, it will ask user to enter **"yes"** if they wish to create samples for animal category and **"no"** for background category. Please note that you would need to run this code multiple times for "yes" and "no" categories as it is better to take sample representation from many videos in your dataset.

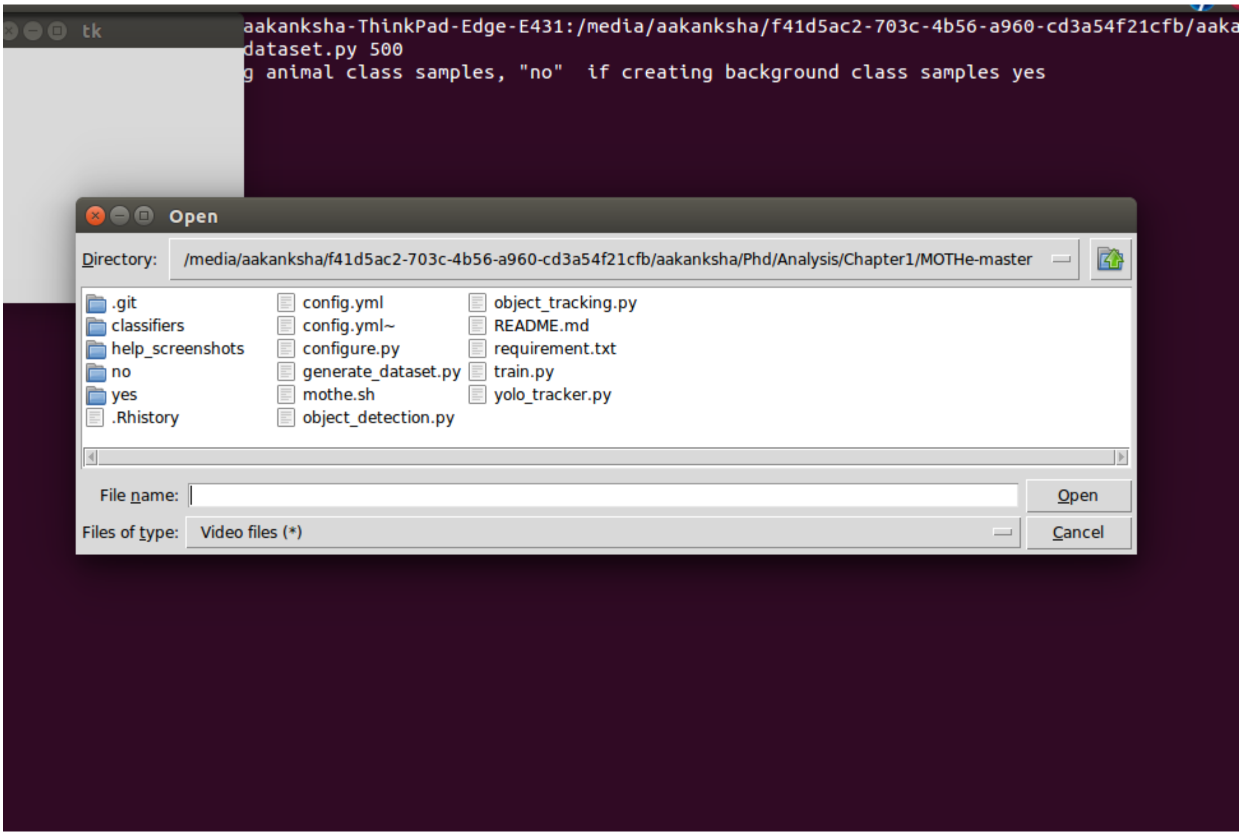


After this input, it will prompt user to select a video from which samples will be saved. After selecting the video, frames from this video will be displayed. User can click on the animals or different points on the background dependingon which category data is being created in the particular execution. Once you have clicked on desired number of points in one frame, press **n** to coninue to the next frame. Press **q** when done and you want to exit.

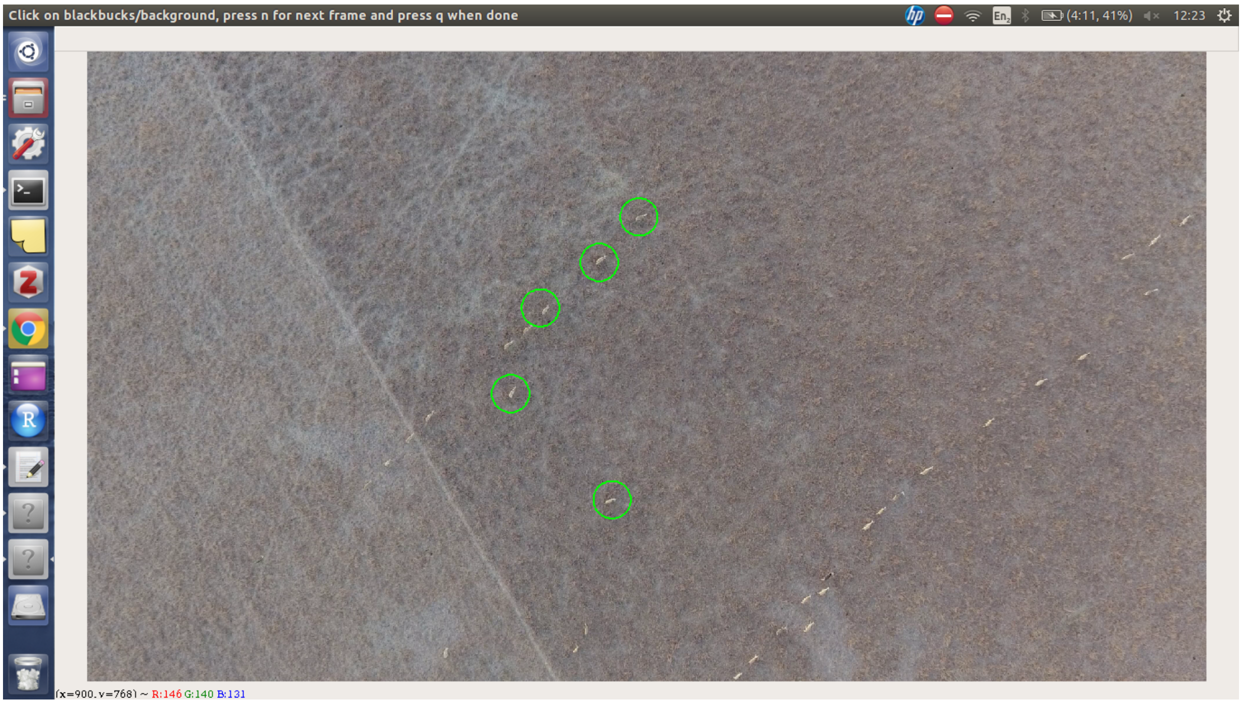


**For testing:** If you wish to test (learn how to run) this module, download our video clips from the [google drive] (). You can then generate samples by choosing any of these videos. If you directly want to proceed to next steps, download our training data from the same drive.

4. **Step 3 - Network training**: Filename - *train.py*

To train the neural network, run the following code in the terminal. The following code trains the neural network on default parameters. If you wish to change the parameters you can do that directly in the script file.

```
$ python train.py
```

You will see the status of training in the command line.

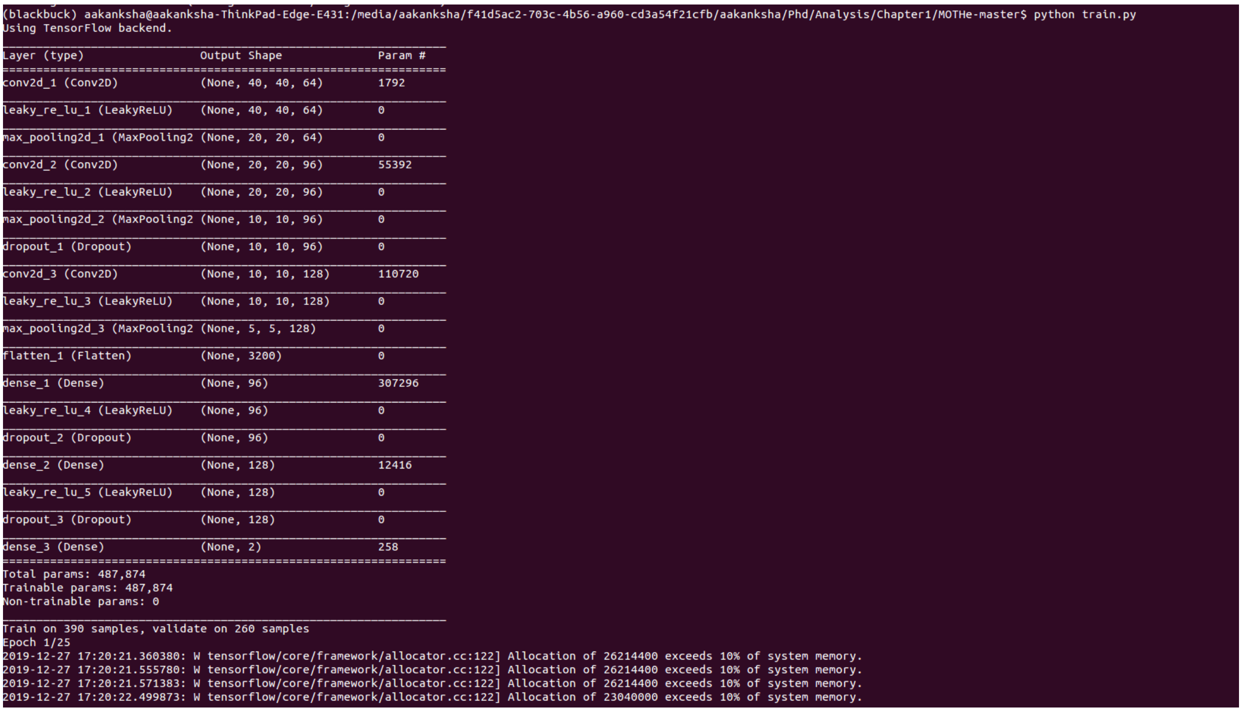

```
(blackbuck) aakanksha@aakanksha-ThinkPad-Edge-E431:/media/aakanksha/f41d5ac2-703c-4b56-a960-cd3a54f21cfb/aakanksha/Phd/Analysis/Chapter1/MOTHe-master$ python train.py
Using TensorFlow backend.
_________________________________________________________________
Layer (type)                 Output Shape              Param #
=================================================================
conv2d_1 (Conv2D)            (None, 40, 40, 64)        1792
_________________________________________________________________
leaky_re_lu_1 (LeakyReLU)    (None, 40, 40, 64)        0
_________________________________________________________________
max_pooling2d_1 (MaxPooling2 (None, 20, 20, 64)        0
_________________________________________________________________
conv2d_2 (Conv2D)            (None, 20, 20, 96)        55392
_________________________________________________________________
leaky_re_lu_2 (LeakyReLU)    (None, 20, 20, 96)        0
_________________________________________________________________
max_pooling2d_2 (MaxPooling2 (None, 10, 10, 96)        0
_________________________________________________________________
dropout_1 (Dropout)          (None, 10, 10, 96)        0
_________________________________________________________________
conv2d_3 (Conv2D)            (None, 10, 10, 128)       110720
_________________________________________________________________
leaky_re_lu_3 (LeakyReLU)    (None, 10, 10, 128)       0
_________________________________________________________________
max_pooling2d_3 (MaxPooling2 (None, 5, 5, 128)         0
_________________________________________________________________
flatten_1 (Flatten)          (None, 3200)              0
_________________________________________________________________
dense_1 (Dense)              (None, 96)                307296
_________________________________________________________________
leaky_re_lu_4 (LeakyReLU)    (None, 96)                0
_________________________________________________________________
dropout_2 (Dropout)          (None, 96)                0
_________________________________________________________________
dense_2 (Dense)              (None, 128)               12416
_________________________________________________________________
leaky_re_lu_5 (LeakyReLU)    (None, 128)               0
_________________________________________________________________
dropout_3 (Dropout)          (None, 128)               0
_________________________________________________________________
dense_3 (Dense)              (None, 2)                 258
=================================================================
Total params: 487,874
Trainable params: 487,874
Non-trainable params: 0
_________________________________________________________________
Train on 390 samples, validate on 260 samples
Epoch 1/25
2019-12-27 17:20:21.360380: W tensorflow/core/framework/allocator.cc:122] Allocation of 26214400 exceeds 10% of system memory.
2019-12-27 17:20:21.555780: W tensorflow/core/framework/allocator.cc:122] Allocation of 26214400 exceeds 10% of system memory.
2019-12-27 17:20:21.571383: W tensorflow/core/framework/allocator.cc:122] Allocation of 26214400 exceeds 10% of system memory.
2019-12-27 17:20:22.499873: W tensorflow/core/framework/allocator.cc:122] Allocation of 23040000 exceeds 10% of system memory.
```

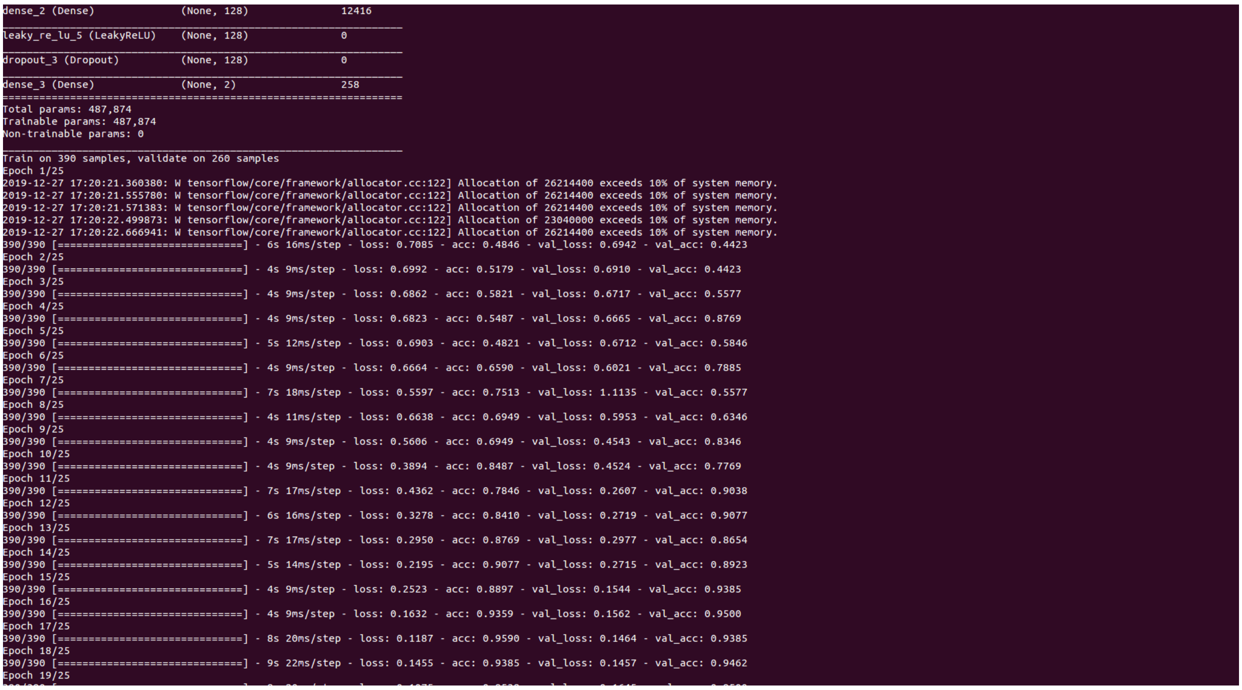

```
dense_2 (Dense)              (None, 128)               12416
_________________________________________________________________
leaky_re_lu_5 (LeakyReLU)    (None, 128)               0
_________________________________________________________________
dropout_3 (Dropout)          (None, 128)               0
_________________________________________________________________
dense_3 (Dense)              (None, 2)                 258
=================================================================
Total params: 487,874
Trainable params: 487,874
Non-trainable params: 0
_________________________________________________________________
Train on 390 samples, validate on 260 samples
Epoch 1/25
2019-12-27 17:20:21.360380: W tensorflow/core/framework/allocator.cc:122] Allocation of 26214400 exceeds 10% of system memory.
2019-12-27 17:20:21.555780: W tensorflow/core/framework/allocator.cc:122] Allocation of 26214400 exceeds 10% of system memory.
2019-12-27 17:20:21.571383: W tensorflow/core/framework/allocator.cc:122] Allocation of 26214400 exceeds 10% of system memory.
2019-12-27 17:20:22.499873: W tensorflow/core/framework/allocator.cc:122] Allocation of 23040000 exceeds 10% of system memory.
2019-12-27 17:20:22.666941: W tensorflow/core/framework/allocator.cc:122] Allocation of 26214400 exceeds 10% of system memory.
390/390 [==============================] - 6s 16ms/step - loss: 0.7085 - acc: 0.4846 - val_loss: 0.6942 - val_acc: 0.4423
Epoch 2/25
390/390 [==============================] - 4s 9ms/step - loss: 0.6992 - acc: 0.5179 - val_loss: 0.6910 - val_acc: 0.4423
Epoch 3/25
390/390 [==============================] - 4s 9ms/step - loss: 0.6862 - acc: 0.5821 - val_loss: 0.6717 - val_acc: 0.5577
Epoch 4/25
390/390 [==============================] - 4s 9ms/step - loss: 0.6823 - acc: 0.5487 - val_loss: 0.6665 - val_acc: 0.8769
Epoch 5/25
390/390 [==============================] - 5s 12ms/step - loss: 0.6903 - acc: 0.4821 - val_loss: 0.6712 - val_acc: 0.5846
Epoch 6/25
390/390 [==============================] - 4s 9ms/step - loss: 0.6664 - acc: 0.6590 - val_loss: 0.6021 - val_acc: 0.7885
Epoch 7/25
390/390 [==============================] - 7s 18ms/step - loss: 0.5597 - acc: 0.7513 - val_loss: 1.1135 - val_acc: 0.5577
Epoch 8/25
390/390 [==============================] - 4s 11ms/step - loss: 0.6638 - acc: 0.6949 - val_loss: 0.5953 - val_acc: 0.6346
Epoch 9/25
390/390 [==============================] - 4s 9ms/step - loss: 0.5606 - acc: 0.6949 - val_loss: 0.4543 - val_acc: 0.8346
Epoch 10/25
390/390 [==============================] - 4s 9ms/step - loss: 0.3894 - acc: 0.8487 - val_loss: 0.4524 - val_acc: 0.7769
Epoch 11/25
390/390 [==============================] - 7s 17ms/step - loss: 0.4362 - acc: 0.7846 - val_loss: 0.2607 - val_acc: 0.9038
Epoch 12/25
390/390 [==============================] - 6s 16ms/step - loss: 0.3278 - acc: 0.8410 - val_loss: 0.2719 - val_acc: 0.9077
Epoch 13/25
390/390 [==============================] - 7s 17ms/step - loss: 0.2950 - acc: 0.8769 - val_loss: 0.2977 - val_acc: 0.8654
Epoch 14/25
390/390 [==============================] - 5s 14ms/step - loss: 0.2195 - acc: 0.9077 - val_loss: 0.2715 - val_acc: 0.8923
Epoch 15/25
390/390 [==============================] - 4s 9ms/step - loss: 0.2523 - acc: 0.8897 - val_loss: 0.1544 - val_acc: 0.9385
Epoch 16/25
390/390 [==============================] - 4s 9ms/step - loss: 0.1632 - acc: 0.9359 - val_loss: 0.1562 - val_acc: 0.9500
Epoch 17/25
390/390 [==============================] - 8s 20ms/step - loss: 0.1187 - acc: 0.9590 - val_loss: 0.1464 - val_acc: 0.9385
Epoch 18/25
390/390 [==============================] - 9s 22ms/step - loss: 0.1455 - acc: 0.9385 - val_loss: 0.1457 - val_acc: 0.9462
Epoch 19/25
```

Once the training is complete, training accuracy will be displayed and trained model will saved with name *mothe_model.h5py* in the folder *classifiers*.

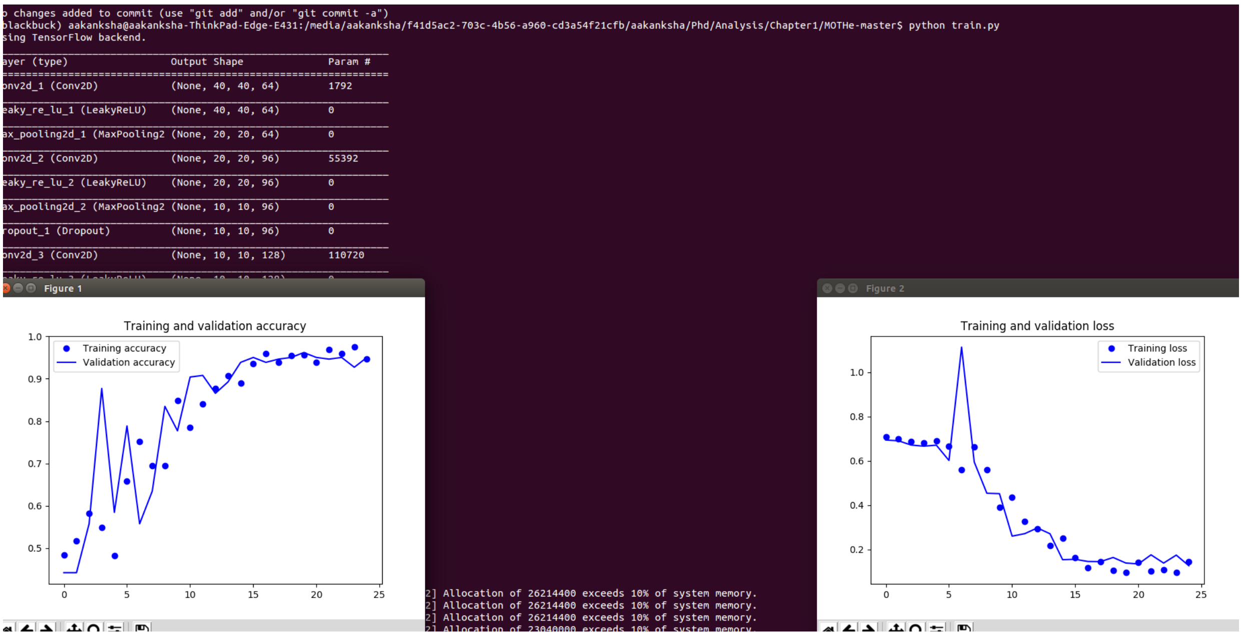


Training accuracy should be at least 98% for the model to work decently, if you wish to increase the accuracy, try training with more data.

**For testing -** You can completely skip this step if you want to run MOTHe on wasp or blackbuck videos. For these videos, trained models are already saved in the repository.

4. **Step 4 - Object detection**: Filename - *object_detection.py*

This module will detect the animals (object of interest) in the video frames. As mentioned earlier, this process is done in two steps - first the code predicts the areas in which animal may be potentially present (localisation) and then these areas are passes to the network for classification task. For localisation, we need thrsholding approach which gives us regions which have animals as well as background noise. To perform this task, user needs to provide threshold values. See section **Parametrization** for details of how to choose threshold values for new datasets.

For object detection, run the following code in the terminal-

```
$ python object_detection.py
```

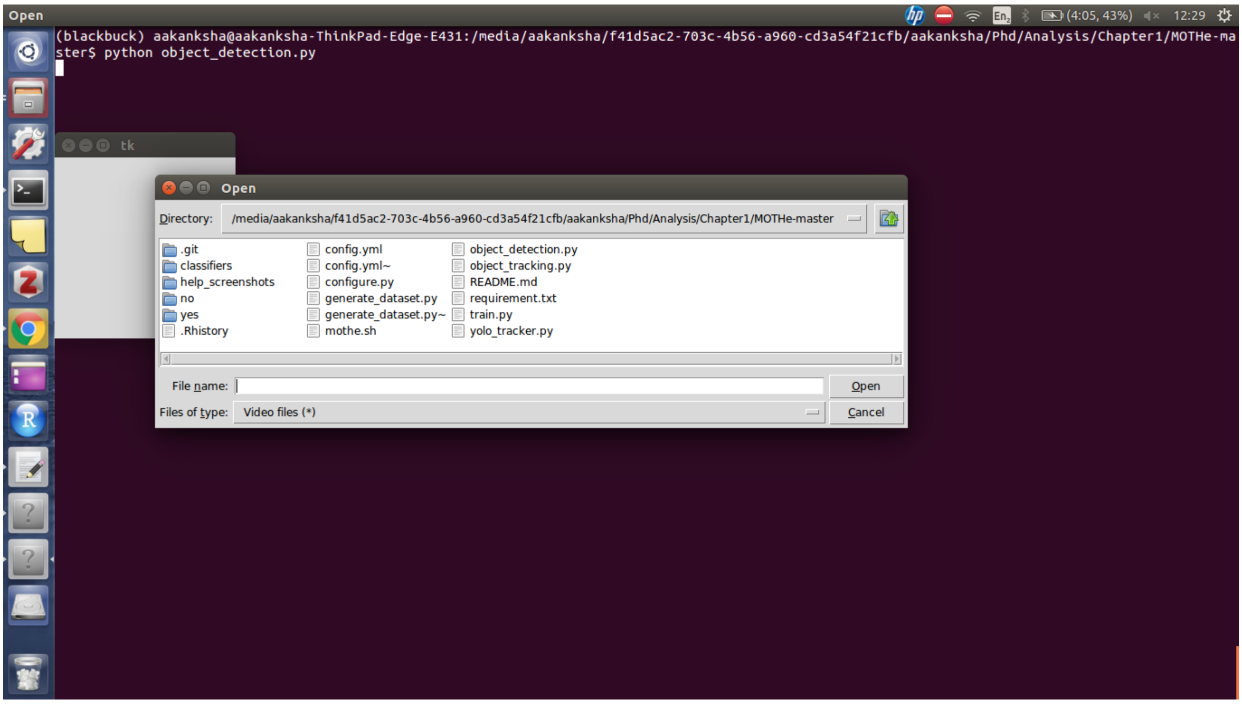


This code will also prompt for number of frames to skip. This will depend on the temporal resolution needed by user. If you need to track animal in each frame then select one, if you want data only 5 frames per second then enter 6. We recommend to not exceed this step size above 6 if user needs to track the individuals. For count data, this number can be as high as number of frames in the video.

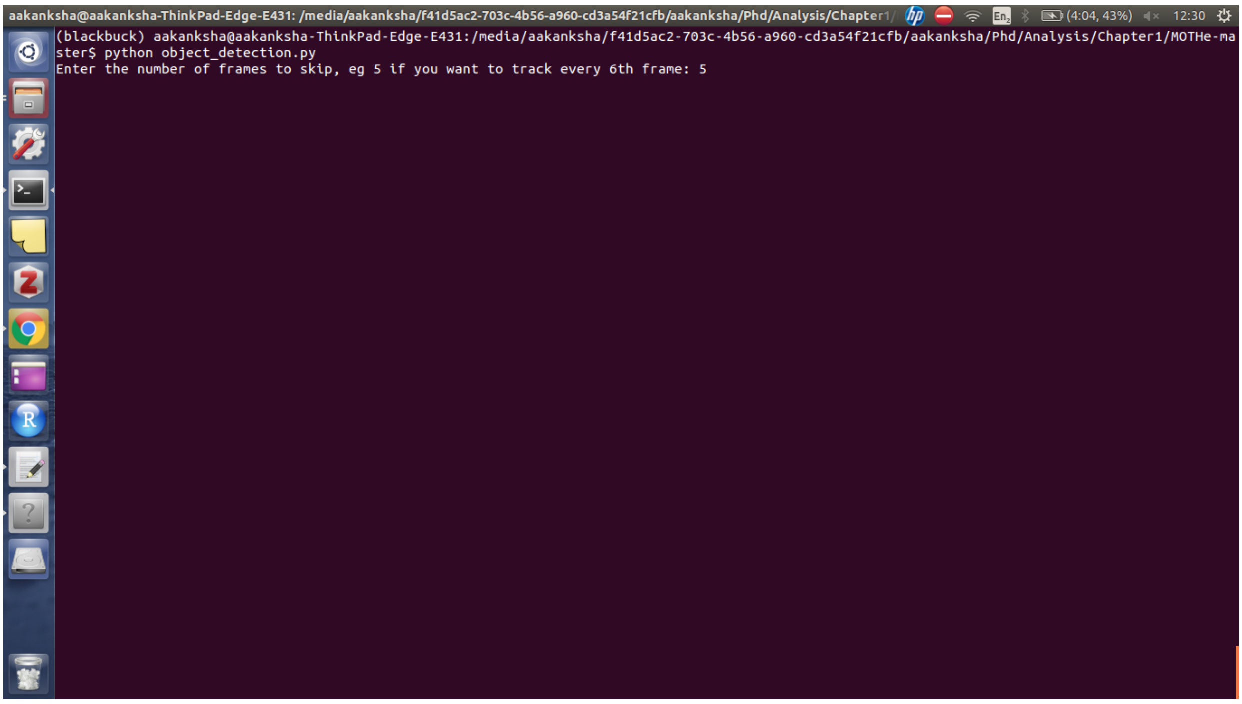


The status and number of frames processed will be shown on the screen-

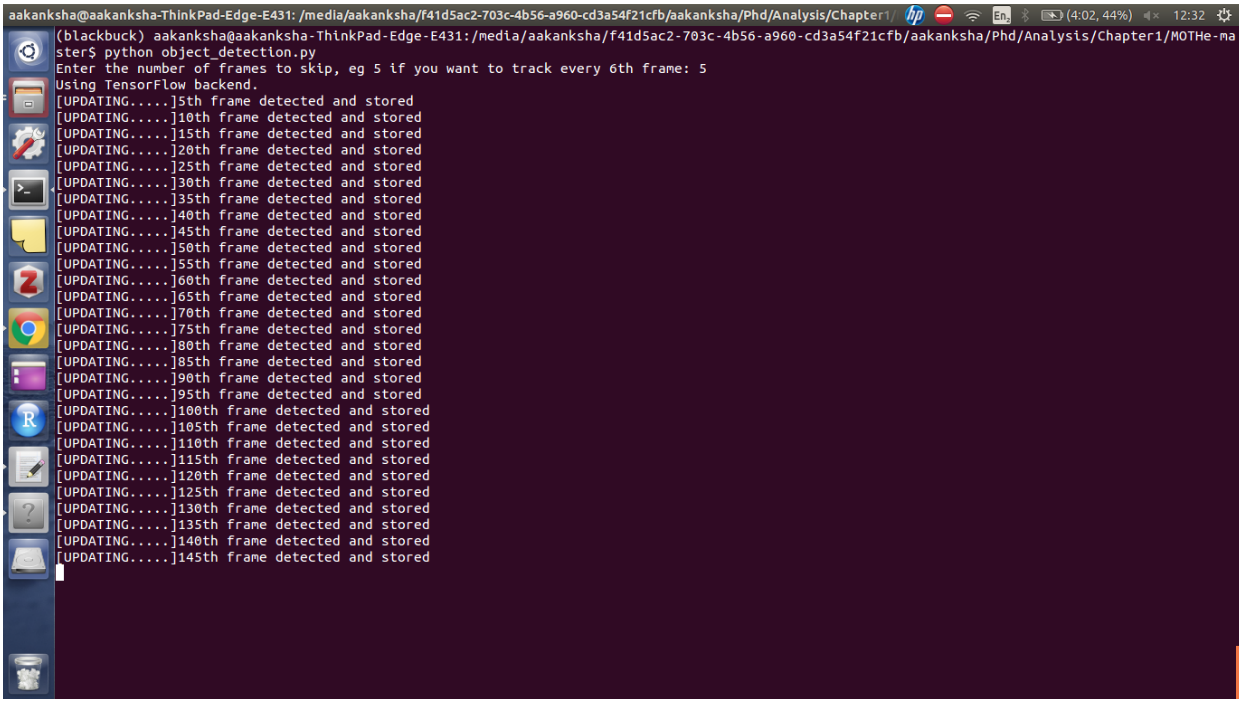


The output from this step is saved in the form of *.csv* files and videos.

**For testing -**

If you wish to test this module on blackbuck or wasp data, you can use default parameters provided in *config.yml*. Open *config.yml* in any text editor and set the values to wasp or blackbuck default mentioned in the comments of these parameters in the file.

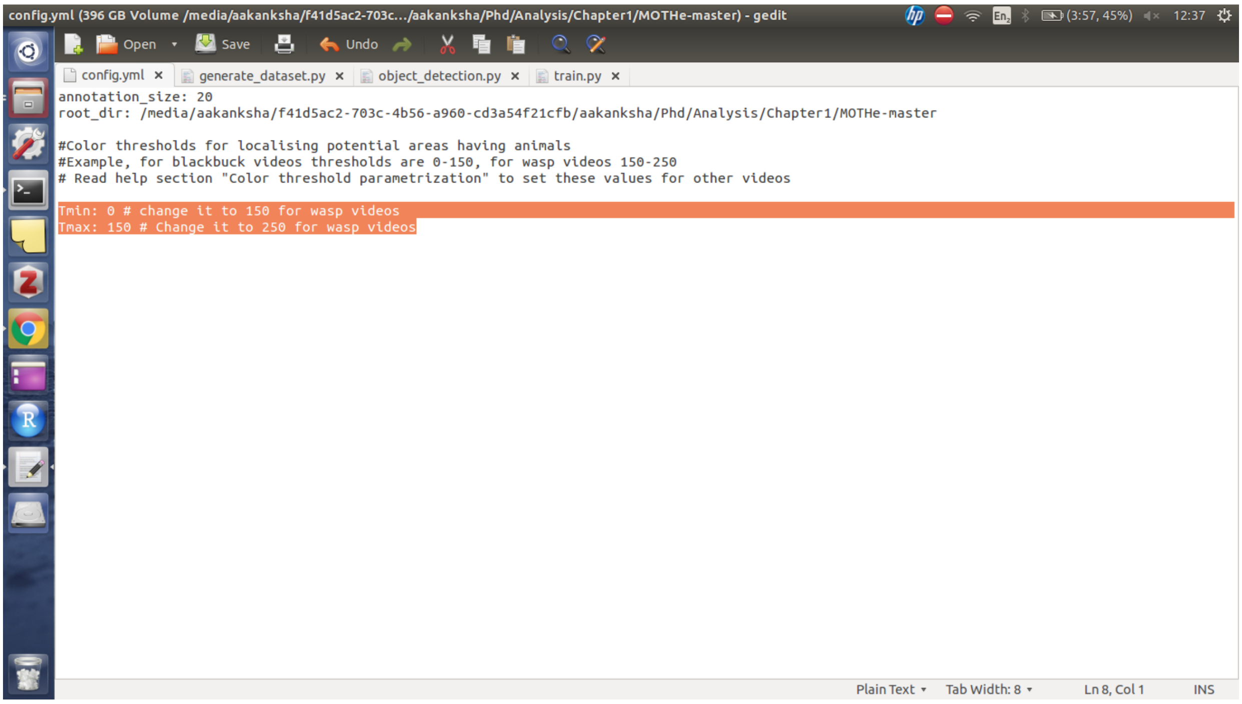


5. **Step 5 - Track linking**: Filename - *object_tracking.py*

This step is used to ascribe unique IDs to the detected animals and it gives us thetrajectoris of the animals. It will use the detections from previous step. Hence, the input for this step would be original video clip and *.csv* generated in the previous step.

Run command -

```
$ python object_tracking.py
```

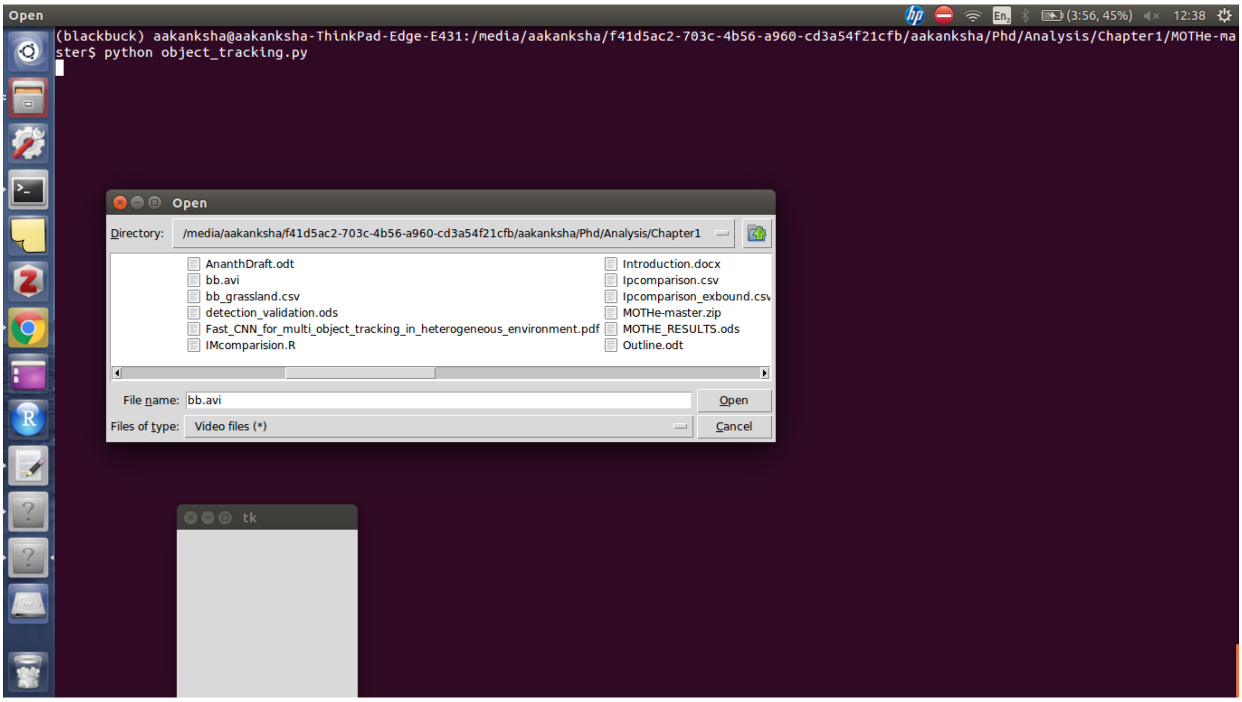


#### HOW MUCH TRAINING DATA DO I NEED?

MOTHe uses a CNN which uses a set of labelled examples to learn the features of the objects. Neural Networks generally work well with huge number of training samples. We recommend using at least 8-10k image examples for the animal category. This number may need to be increased if the animal of interest shows a lot of variation in morphology. For example, if males and females are of different colors, it is important to include sufficient examples for both of them. Similarly, if the background is too heterogeneous then you may need more training data (around 1500-2000 samples for different types of variations in the background). For example to train the MOTHe on our blackbuck videos, we used 9800 cropped samples for blackbuck (including males and females) and 19000 samples for the background because background included grass, soil, rocks, bushes, water etc.

#### CHOOSING COLOR THRESHOLDS

The object detection steps requires user to enter threshold values in the config files. Object detection in MOTHe works in two steps, it first uses a color filter to identify the regions in the image on which to run the classification. We use color threshold to select these regions. You can see the values of thresholds for blackbuck and wasp videos in the _config.yml_file. If you are running MOTHe on a new dataset, there are two ways to select appropriate threshold values:

1. You may open some frames from different videos in an interactive viewer (for example MATLAB image display), you can then click on the pixels on animal and check the RGB values (take the avergae of all 3 channels). Looking at these values in multiple instances will give you an idea to choose a starting minimum and maximum threshold. Run the detection with these thresholds and you can improve the detection by hit and trial method to tweak the threshold.

2. You can compare your videos to wasp and blackbuck videos and start with threshold values to which your data is more similar. For example, if your animal looks more similar to blackbuck in color and lighting conditions, you may start with default thresholds and improve the detection by changing lower and upper threshold by little amount at a time.

======= Software to detect and track animals in their natural environment
